# Grad-seq analysis of *Enterococcus faecalis* and *Enterococcus faecium* provides a global view of RNA and protein complexes in these two opportunistic pathogens

**DOI:** 10.1101/2022.11.09.515799

**Authors:** Charlotte Michaux, Milan Gerovac, Elisabeth E. Hansen, Lars Barquist, Jörg Vogel

**Affiliations:** Institute for Molecular Infection Biology, University of Würzburg, Würzburg, Germany; Helmholtz Institute for RNA-based Infection Research (HIRI), Helmholtz Centre for Infection Research (HZI), Würzburg, Germany; Faculty of Medicine, University of Würzburg, Würzburg, Germany

**Keywords:** Grad-seq, *E. faecalis*, *E. faecium*, RNA-binding protein, KhpA, KhpB, KH domain, sRNA, tRNA, 3’UTR

## Abstract

*Enterococcus faecalis* and *Enterococcus faecium* are major nosocomial pathogens. Despite their relevance to public health and their role in the development of bacterial antibiotic resistance, relatively little is known about gene regulation in these species. RNA–protein complexes serve crucial functions in all cellular processes associated with gene expression, including post-transcriptional control mediated by small regulatory RNAs (sRNAs). Here, we present a new resource for the study of enterococcal RNA biology, employing the Grad-seq technique to comprehensively predict complexes formed by RNA and proteins in *E. faecalis* V583 and *E. faecium* AUS0004. Analysis of the generated global RNA and protein sedimentation profiles led to the identification of RNA-protein complexes and putative novel sRNAs. Validating our data sets, we observe well-established cellular RNA-protein complexes such as the 6S RNA-RNA polymerase complex, suggesting that 6S RNA-mediated global control of transcription is conserved in enterococci. Focusing on the largely uncharacterized RNA-binding protein KhpB, we use the RIP-seq technique to predict that KhpB interacts with sRNAs, tRNAs, and untranslated regions of mRNAs, and might be involved in the processing of specific tRNAs. Collectively, these datasets provide departure points for in-depth studies of the cellular interactome of enterococci that should facilitate functional discovery in these and related Gram-positive species. Our data are available to the community through a user-friendly Grad-seq browser that allows interactive searches of the sedimentation profiles (https://resources.helmholtz-hiri.de/gradseqef/).

## INTRODUCTION

Enterococci are ubiquitious Gram-positive members of the gut microbiome. Once considered harmless commensals, they have recently been re-assigned as nosocomial pathogens because the two Enterococcus species *Enterococcus faecalis* and *Enterococcus faecium* were found to cause life-threatening infections (Hidron et al., 2008; Kristich et al., 2014; Weiner et al., 2016). Enterococci often carry intrinsic or acquired resistance to a wide range of antimicrobial agents, which limits treatment options (Hollenbeck and Rice, 2012; Kristich et al., 2014; Mundy et al., 2000). Importantly, they have also been shown to transmit antibiotic resistance genes to other Gram-positive and even Gram-negative species (Courvalin, 1994; Weigel et al., 2003).

Despite the clinical relevance of *E. faecalis* and *E. faecium* (Arias and Murray, 2012; Fiore et al., 2019; Kristich et al., 2014; Van Tyne and Gilmore, 2014), general gene regulatory mechanisms in these bacteria remain poorly understood (DebRoy et al., 2014; Weaver, 2019). Given the central function of RNA-protein complexes in gene regulation, the global categorization of RNA-binding proteins (RBPs) and their RNA partners provides an important framework to address this question (Gerovac et al., 2021a; Holmqvist and Vogel, 2018). RBPs are also known to be important co-factors during post-transcriptional regulation mediated by small noncoding RNAs (sRNAs). Most sRNAs act to repress target mRNAs at the level of translation or affect stability by short base pairing interactions that also involve an sRNA chaperone, as exemplified by most Hfq- and some ProQ-dependent sRNAs in Gram-negative bacteria (Holmqvist et al., 2020; Hör et al., 2020c; Ponath et al., 2022; Quendera et al., 2020). While the physiological importance of this mode of gene regulation is well-established in Gram-negative bacteria, it is still unclear if a similarily broad mechanism exists in Gram-positive bacteria. Indeed, although sRNAs have been identified in Gram-positive bacterial species, including *E. faecalis* and *E. faecium* (Fouquier d’Hérouel et al., 2011; Innocenti et al., 2015; Michaux et al., 2020; Shioya et al., 2011; Sinel et al., 2017), an RBP with a global function comparable to the sRNA chaperones Hfq, ProQ, and CsrA present in Gram-negative bacteria has not yet been discovered. More generally, the discovery of RBPs with gene regulatory functions has been hampered by the absence of experimental global data sets to predict the molecular complexes in which transcripts and proteins of enterococci engage.

Grad-seq is a recently developed approach to discover RBPs and to determine native RNA-protein and protein-protein complexes (Smirnov et al., 2016). The method is based on the separation of soluble cellular complexes by a classical glycerol gradient, followed by high-throughput RNA sequencing (RNA-seq) and mass spectrometry (MS) analyses of the individual gradient fractions. Potential RBPs are then predicted by searching for correlations between in-gradient behavior of cellular proteins and transcripts. Thus far, Grad-seq has led to the identification of (i) ProQ as a global sRNA-binding protein in *Salmonella enterica* serovar Typhimurium (Smirnov et al., 2016), (ii) new components of well-established complexes, such as the broadly conserved protein YggL as a factor associated with the ribosome in *Escherichia coli* (Hör et al., 2020a), (iii) a new mechanism of exonucleolytic sRNA activation in *Streptococcus pneumoniae* (Hör et al., 2020b), (iv) the RNA complexome of *Pseudomonas aeruginosa* under bacteriophage infection (Gerovac et al., 2021b) and (v) a new RBP in *Clostridium difficile* (Lamm-Schmidt et al., 2021). In addition, GradR as a new variant of Grad-seq that includes an RNase treatment step helped to discover a hitherto unrecognized, plasmid-encoded FinO-domain RBP in *Salmonella* (Gerovac et al., 2020).

Here, we applied Grad-seq to chart the landscape of RNA-protein and protein-protein complexes in *E. faecalis* and *E. faecium*. Our analysis of the sedimentation profiles reveals previously hidden sRNAs and predicts new candidate enterococcal RBPs. One of these is the conserved KH domain protein KhpB, which was recently shown to interact with RNAs in *S. pneumoniae* and *C. difficile* (Hör et al., 2020b; Lamm-Schmidt et al., 2021; Riediger et al., 2021; Zheng et al., 2017). KH domains have been found in bacterial RBPs with a wide range of functions, such as the enzymes polynucleotide phosphorylase (PNPase) and RNase Y or the transcription elongation factor NusA (Olejniczak et al., 2021). In the present work, a pilot analysis of cellular RNA ligands of *E. faecalis* KhpB suggests that its RNA interactome is mainly composed of sRNAs, tRNAs and untranslated regions (UTRs), and that this RBP might have an RNA processing function. Our Grad-seq data sets, available in a user-friendly browser (https://resources.helmholtz-hiri.de/gradseqef/), provide a global resource to better understand transcriptional and post-transcriptional regulation in *E. faecalis* and *E. faecium*, the two pathogenic species of the *Enterococcus* family.

## RESULTS

### Grad-seq recapitulates known RNA-protein complexes in E. faecalis and E. faecium

To establish an atlas of cellular protein-protein and protein-RNA complexes in *E. faecalis* and *E. faecium*, we performed Grad-seq (see Fig. 1A for general workflow) on bacterial lysates obtained at late exponential phase (OD_600_=2) in M17 rich media. We chose this growth phase based on a previous study (Michaux et al., 2020) in which we had mapped transcription start sites (TSSs) in *E. faecalis* and *E. faecium*, and which showed that the vast majority of genes are expressed in this condition. Whole-cell lysates were analysed in a glycerol gradient by ultracentrifugation and subsequently fractionated in 20 fractions plus the pellet. The absorbance of each fraction at 260 nm revealed a characteristic global profile (Fig. 1B for *E. faecalis*; Fig. S1A for *E. faecium*). Specifically, the bulk peak at the top of the gradient represents low-molecular weight (LMW) fractions in which free proteins and small complexes accumulate. In the high-molecular weight (HMW) fractions, two characteristic peaks around fractions 10 and 15 are observed, which correspond to the small (30S) and large (50S) ribosomal subunits, respectively; this pattern is very similar to the ribosome peaks observed in the closely related Gram-positive species, *S. pneumoniae* (Hör et al., 2020b).

**Figure 1.**
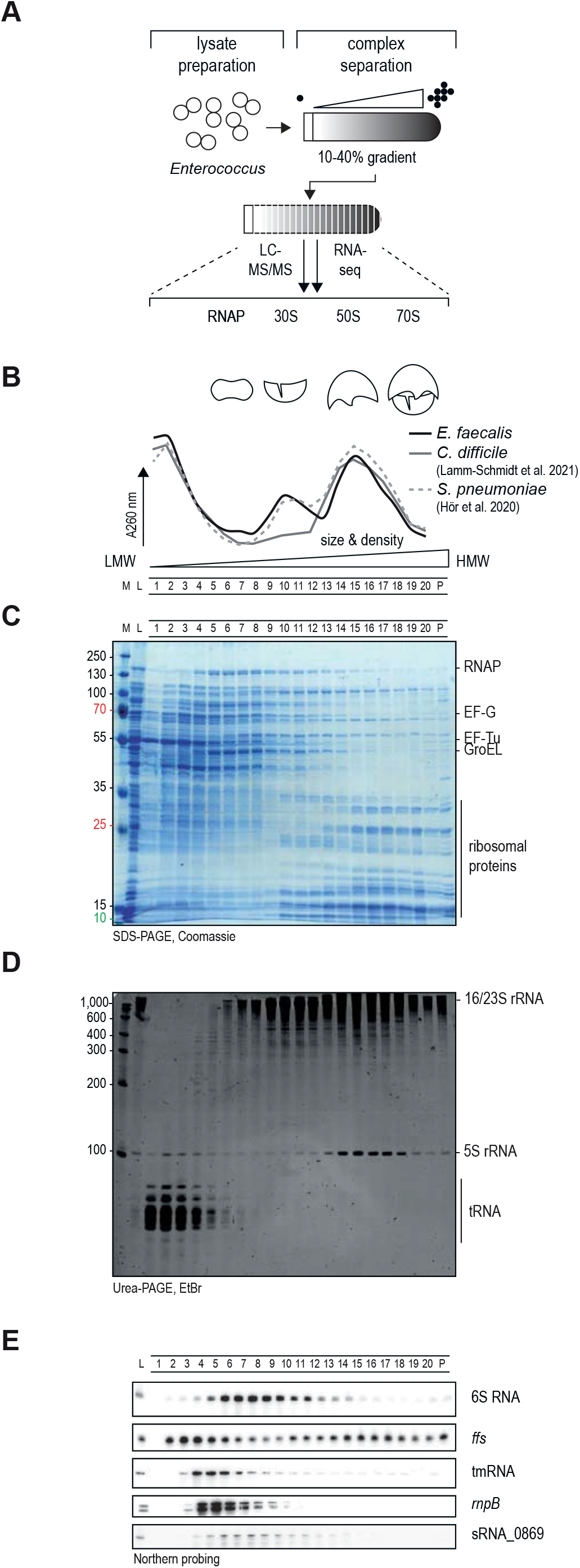
Grad-seq in *E. faecalis*. (A) General workflow: cellular lysate was sedimented in a glycerol gradient, fractionated and analyzed by RNA-seq or mass-spectrometry to reveal sedimentation profiles. (B) The 260 nm absorption profile shows a bulk signal in LMW fractions followed by a peak for the small 30S ribosomal subunit and a peak for the large 50S subunit. This profile is comparable to profiles obtained previously for *S. pneumoniae* and *C. difficile*. (C) Fractions were run on an SDS-PAGE and abundant proteins were detected by Coomassie staining. (D) Fractions were run on a Urea-PAGE and abundant RNAs were detected by ethidium bromide staining. (E) Northern blots were probed for the indicated RNAs to assess their sedimentation profile.

Total protein and RNA were extracted for each fraction and analyzed by SDS-PAGE or urea-PAGE to visualize abundant proteins (Figs. 1C, S1B) and transcripts (Figs. 1D, S1C), respectively. Judging by abundance and size, we observed ribosomal proteins in the HMW fractions, as expected from the absorbance profiles (Fig. 1B-C). Nonetheless, the most abundant proteins in both species were present in the LMW fractions, such as elongation factors EF-Tu (43 kDa) and EF-G (76 kDa) in fractions 2 to 5, the glycerol kinase GlpK (55 kDa) in fractions 1 to 4, or the complex of the chaperone GroEL (60 kDa) in fractions 7 to 13 (Figs. 1C, S1B). Subunits of the RNA polymerase (RNAP) sedimented in fractions 4 to 8 (Figs. 1C, S1B). Regarding transcripts (Figs. 1D, S1C), highly abundant tRNAs sedimented in LMW fractions 1 to 4, whereas the 16S and 23S/5S ribosomal RNAs (rRNAs) occurred in the ribosomal fractions, as expected.

For additional quality evaluation, we performed independent northern blot analysis of some well-characterized stable transcripts, such as 6S RNA, 4.5S RNA (*ffs* gene), tmRNA, M1 RNA (*rnpB*) or sRNA_0869, an sRNA previously identified in *E. faecalis* (Fouquier d’Hérouel et al., 2011; Shioya et al., 2011) (Figs. 1E, S1D). Where comparable, these sedimentation profiles resemble reported profiles in *E. coli*, *Salmonella*, *S. pneumoniae*, *P. aeruginosa* and *C. difficile* (Gerovac et al., 2021b; Hör et al., 2020a, 2020b; Lamm-Schmidt et al., 2021; Smirnov et al., 2016). For example, tmRNA, which forms a ribonucleoprotein complex with the protein SmpB that rescues stalled ribosomes (Huter et al., 2017) sedimented in fractions 3 to 8, as expected by a predicted complex size of 32 kDa (Figs. 1E, S1D). We observed 6S RNA in fractions 5 to 9, corresponding to the distribution of the protein subunits of RNAP (Figs. 1E, S1D); this suggests that enterococci use a 6S RNA-dependent mechanism for global control of transcription, as reported in several other species (Wassarman, 2018). M1 RNA, the catalytic RNA subunit of RNaseP (Frank and Pace, 1998), appeared in fractions 3 to 7, as previously seen in *Streptococcus* (Hör et al., 2020b) (Figs. 1E, S1D).

The fractionated gradient was subjected to high-throughput RNA sequencing and mass spectrometry in order to compile sedimentation profiles for proteins and RNAs and enable global predictions of macromolecular complexes. Protein and RNA abundance was normalized via an external spike-in (Tables S1, S2). To benchmark the quality of the global dataset, high-throughput sedimentation profiles of well-established complexes were evaluated for co-sedimentation (Fig. 2). For example, the RNA profiles of 6S, 4.5S RNA, and rRNAs showed excellent correlation with the profiles of proteins with which they are known to form complexes with—the RNAP subunits, the Ffh protein, and the ribosomal proteins, respectively (Figs. 2A, S2A). Altogether, the high correlation between the profiles of proteins and RNAs known to form complexes encouraged us to further analyze the data in a high-throughput fashion.

**Figure 2.**
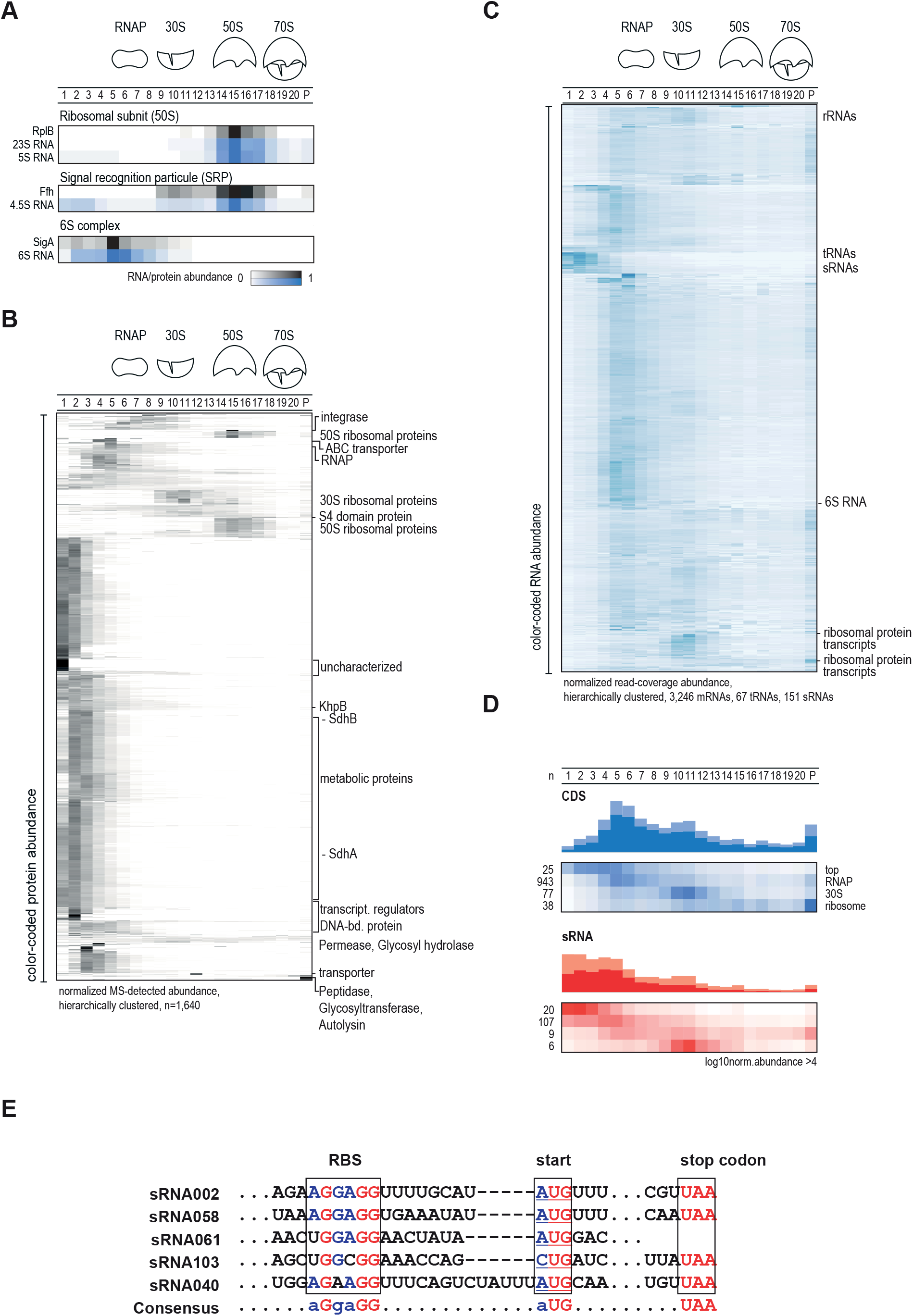
Grad-seq sedimentation profiles for *E. faecalis*. (A) Sedimentation profiles of known RNA-protein complexes, specifically the ribosomal 50S subunit complex, the signal recognition particle (SRP) and the 6S complex (an association between 6S and the sigma factor SigA in RNAP fractions). (B) Heatmaps based on unsupervised clustering of the sedimentation profiles of *E. faecalis* proteins. Clusters that indicate known or new complexes have been highlighted. (C) Heatmaps based on unsupervised clustering of the sedimentation profiles of *E. faecalis* RNA. RNA sedimentation profile patterns can be detected, for example tRNA clusters, sRNAs located in the LMW fractions, or ribosomal RNA in the HMW fractions. (D) Averaged profiles of CDSs and sRNAs. (E) Partial alignment of the 30S-associated sRNAs. The putative RBS, start and stop codons are framed.

### Grad-seq recovers the majority of E. faecalis and E. faecium proteins and transcripts

We used unsupervised clustering of the sedimentation profiles of *E. faecalis* (Fig. 2B-C) and *E. faecium* (Fig. S2B-C) to generate heatmaps. For *E. faecalis*, a total of 1,640 proteins were detected, representing >50% of the 3,240 annotated proteins. The corresponding heat-map (Fig. 2B) highlights different clusters of proteins that share a similar sedimentation profile, including ribosomal proteins or metabolic enzymes, such as the succinyl deshydrogenase complex composed of ShdA and SdhB with a MW of 55kDa. For *E. faecium*, a total of 1,654 proteins were detected, representing >58% of the 2,826 total annotated proteins (Fig. S2).

For cellular RNA species, we used our recent dRNA-seq based single-nucleotide transcriptome annotation in *E. faecalis* (Michaux et al., 2020), and recovered in-gradient distributions of 3,246 mRNAs, 67 tRNAs, 151 sRNAs, 291 3’UTRs and 1,456 5’UTRs (Fig. 2C). For *E. faecium*, 3,049 mRNAs, 47 tRNAs, 129 sRNAs, 342 3’UTRs and 1,470 5’UTRs were detected (Fig. S2C). The fact that we detected more mRNAs than the total number of annotated proteins in both species can be explained by the incomplete annotation of *E. faecalis* and *E. faecium* proteins; currently, not every CDS is present in the Uniprot database. Overall, the total number of transcripts and proteins for which we could establish sedimentation profiles via Grad-Seq in *E. faecalis* and *E. faecium* is comparable to previous Grad-seq experiments in others species, such as *C. difficile*, *E. coli* or *Streptococcus pneumoniae* (Hör et al., 2020b, 2020a; Lamm-Schmidt et al., 2021).

### RNA sedimentation profiles reveal new UTR-derived sRNA candidates

Global clustering of RNA sedimentation profiles allows the identification of groups of mRNAs or sRNAs that might be part of distinct RNA-protein complexes. While in both species some mRNAs (here designated as coding sequences, CDS) were found in fractions 4 to 6 (Figs. 2D, S2D), many others peaked in the RNAP fractions, in the ribosomal fractions or were found in the pellet, indicating association with the transcription or translation machineries (Figs. 2D, S2D). Interestingly, we observed a more pronounced sedimentation of mRNAs in HMW fractions in *E. faecium* (Fig. S2D) than we did in *E. faecalis* (Fig. 2D). This suggests that in the late exponential growth phase, *E. faecium* has a more active protein synthesis compared to *E. faecalis*.

Focusing on sRNAs, we detected a cluster in LMW fractions, which we predict to correspond to largely protein-free sRNAs (Figs. 2D, S2D). A second cluster, more visible in *E. faecium* than *E. faecalis*, is present in fractions 9 to 12; i.e., these sRNAs occur in the 30S ribosomal fractions. There are two tempting possible explanations for this sedimentation profile: firstly, these sRNAs might be associated with translated mRNAs as part of their regulatory activity; alternatively, these sRNAs themselves might be translated, because they contain small open-reading frames (ORFs), perhaps with non-canonical start codons that have been overlooked so far. The latter phenomenon has recently been established in *E. coli* through Grad-seq analysis. RyeG, presumed to be a noncoding sRNA, was found to peak in the 30S fractions of the gradient and was shown to contain a previously unnoticed ORF of 48 amino acids (aa) starting with a GUG start codon (Hör et al., 2020a; Weaver et al., 2019). To address this possibility in *E. faecalis* and *E. faecium*, we performed sequence alignments for 6 sRNAs (sRNA_002; sRNA_040; sRNA_053; sRNA_058; sRNA_061; sRNA_103) that are present in the 30S fraction (Fig. 2D). Five of these sRNAs contain a potential Shine-Dalgarno (SD) sequence, followed by a start codon and, for four of them, a stop codon, revealing possible small OFRs for sRNA_002 (15 aa), sRNA_040 (14 aa), sRNA_058 (20 aa) and sRNA_103 (19 aa) (Fig. 2E).

For in-gradient distributions of individual mRNAs, we observed instances of strikingly different sedimentation profiles of annotated CDS compared to their respective UTRs (Figs. 3A and S2E, Table S4). Typically, all regions of an mRNA would be expected to sediment in the same fraction, and this anomaly might therefore indicate the presence of potential 5’ or 3’-UTR-derived sRNAs (Adams and Storz, 2020; Ponath et al., 2022). Searching for such noncanonical in-gradient distributions, we identified 20 and 41 UTR-derived candidate sRNAs for *E. faecalis* and *E. faecium*, respectively (based on the absolute value differences of at least 5 units between the relative position of the CDS and UTR (see Methods for details; Table S4). As an example the hypothetical gene *EF_3018* is shown in Fig. 3B. Its 5’UTR does not co-fractionate with its CDS, indicative of a possible 5’UTR derived sRNA. Additional examples include the 5’UTRs of the polysaccharide lyase *EF_3023* and the small metabolite transporter *EF_0359*. These instances highlight the power of Grad-seq for extended sRNA annotation.

**Figure 3.**
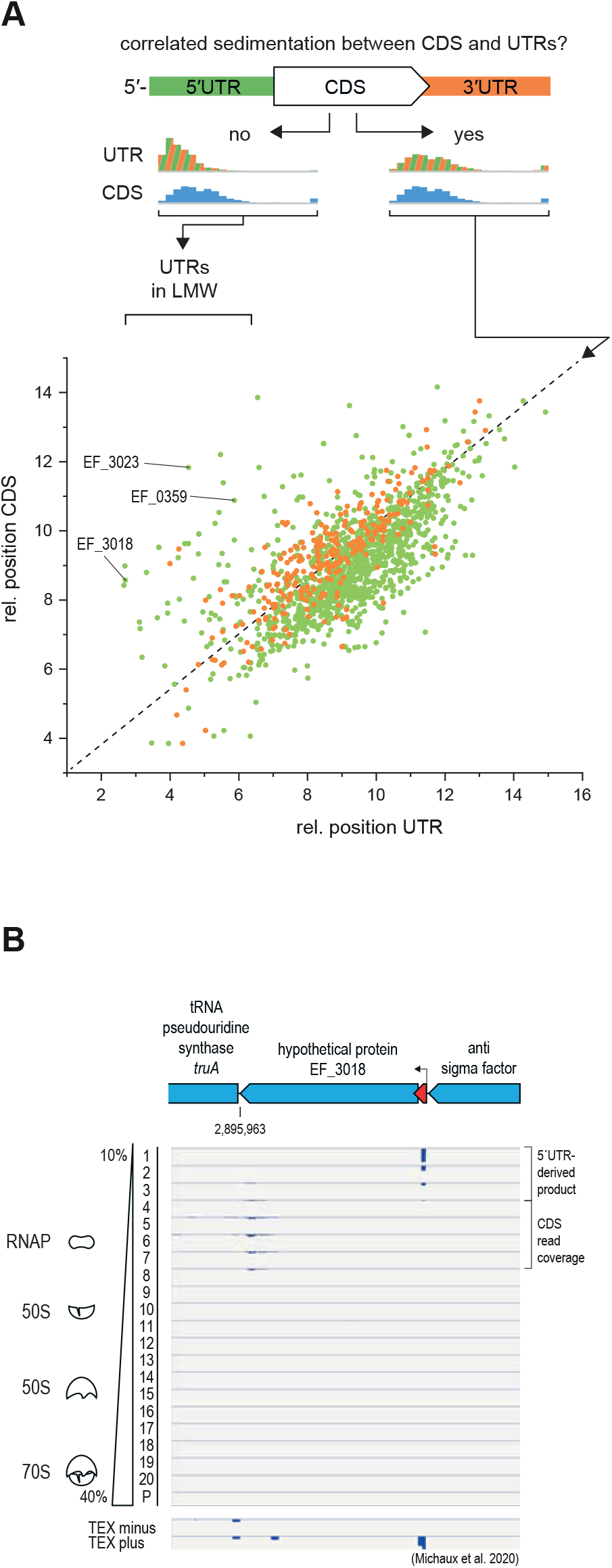
Sedimentation of CDSs and their correspondeding UTRs. (A) Comparison of the relative positions of CDSs and their UTRs in the sedimentation gradient. Outliers indicate a different sedimentation between the UTR and the CDS. (B) Sedimentation profile at nucleotide resolution of EF_3018, one of the outliers labeled in (A).

### Grad-seq identifies new and established protein complexes

Our sedimentation profiles constitute a resource for the identification of molecules that may associate in stable cellular complexes. The strongest indicator of complex formation is the presence of proteins in HMW fractions. Prominent examples include some tRNA synthetases, chaperones such as GroEL, transcription termination factors such as Rho and the subunits of RNAP, as well as numerous metabolic enzymes (Fig. 4A); these proteins often peak in fractions >2 with molecular sizes larger than their free form (>200 kDa). To obtain a high-level view of the sedimentation profiles of cellular proteins, we applied principal-component analysis (PCA) to cosegregate the sedimentation profiles by position and complexity in two components (Fig. 4B for *E. faecalis*, Fig. 4C for *E. faecium*; maximal peak fraction >8 with the principal-component coordinates given in Table S1). As a result, the sedimentation profile complexity is reduced from 21 fractions to two components (Fig. 4B-C). A number of well-characterized complexes aggregate, in accordance with their sedimentation profiles. For instance, in *E. faecalis*, three proteins of the pyruvate dehydrogenase (PDH) complex – PdhA, PdhB and AceF – cluster together, and so did many ribosomal proteins (Fig. 4C).

**Figure 4.**
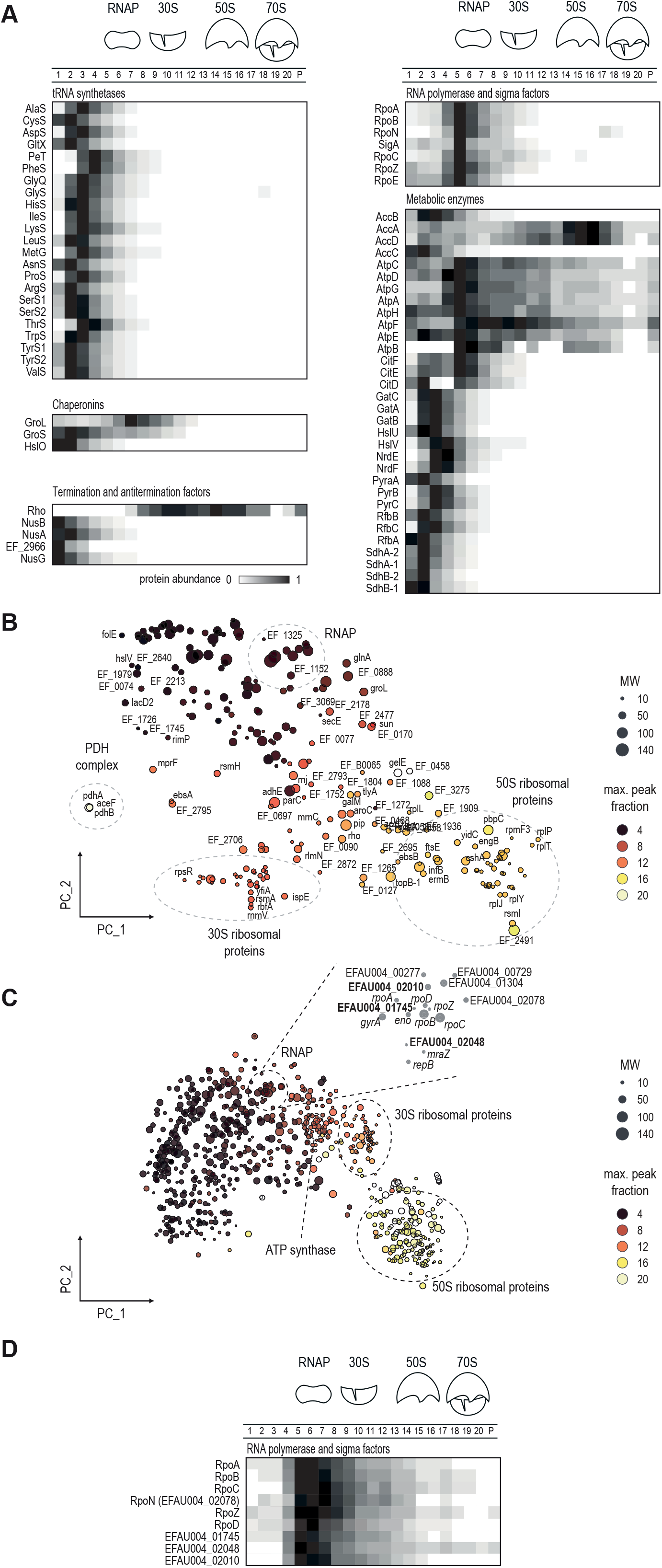
Sedimentation of *E. faecalis* and *E. faecium* protein complexes. (A) Examples of sedimentation profiles of known complexes in *E. faecalis*. (B, C) Clustering of sedimentation profiles by principal component analysis for *E. faecalis* (B) and *E. faecium* (C). Dashed circles indicate protein complex clusters. (D) Sedimentation profiles of the RNAP cluster in *E. faecium*.

This approach is especially valuable for the initial identification of protein complexes in less studied bacterial species. For example, the protein annotation in *E. faecium* lacks many RNAP-associated sigma factors that are described in *E. faecalis*. To identify factors that might form a complex with *E. faecium* RNAP, we zoomed into the PCA plot to explore which of the factors that share similar sedimentation profiles with RNAP might be potential candidates for RNAP interactors (Fig. 4C). As a positive control, we detected EFAU004_02078, which resembles the *E. coli* sigma factor RpoN and is so far the only *E. faecalis* protein annotated in UniProt as a sigma factor. In addition, the hypothetical proteins EFAU004_01745 and EFAU004_02048, and the ABC-transporter EFAU004_02010 shared nearly identical sedimentation profiles with RNAP subunits and lend themselve for further investigation as novel sigma or transcription factors (Fig. 4D). In summary, evaluation of factors that have a highly correlated sedimentation profile with known protein complexes constitutes a first step to predict function for an unknown protein and establish its involvement in a stable complex.

### The emerging RNA-binding protein KhpB is conserved in enterococci

RBPs usually accumulate in the first fractions of the gradient (Gerovac et al., 2020) and can be difficult to discriminate between by PCA due to their small molecular weight and their transient interactions with RNAs. Importantly, previous Grad-seq analysis in Gram-negative bacteria often observed tailing towards the central gradient fractions for established global RBPs such as Hfq and ProQ (Hör et al., 2020a; Smirnov et al., 2016); when the samples were pretreated with RNase, these RBPs would shift towards LMW fractions (Gerovac et al., 2020). Of note, *E. faecalis* and *E. faecium* lack CsrA, Hfq, and ProQ, the three major global RBPs associated with sRNA-mediated gene regulation in Gram-negative bacteria (Christopoulou and Granneman, 2021). Although other RBPs have been reported in Gram-positive bacteria, e.g., diverse members of the cold shock protein (Csp) family, the ppGpp synthetase RelQ or the ribosomal protein S1, there is currently no evidence for a global sRNA-binding protein in *E. faecalis* or *E. faecium* (Christopoulou and Granneman, 2021). Thus, we examined a number of annotated RBPs in these two organisms (Fig. S3). Interestingly, they all showed tailing towards RNAP and ribosomal fractions, which could be indicative of stable interactions with other cellular partners. Interestingly, we also observed that the putative RBPs KhpA and KhpB (Olejniczak et al., 2021) showed strong tailing towards the RNAP and 30S fractions (Fig. 5A). Moreover, their profiles correlate with sRNA clusters (Fig. 2D), making them appealing candidates for further characterization.

**Figure 5.**
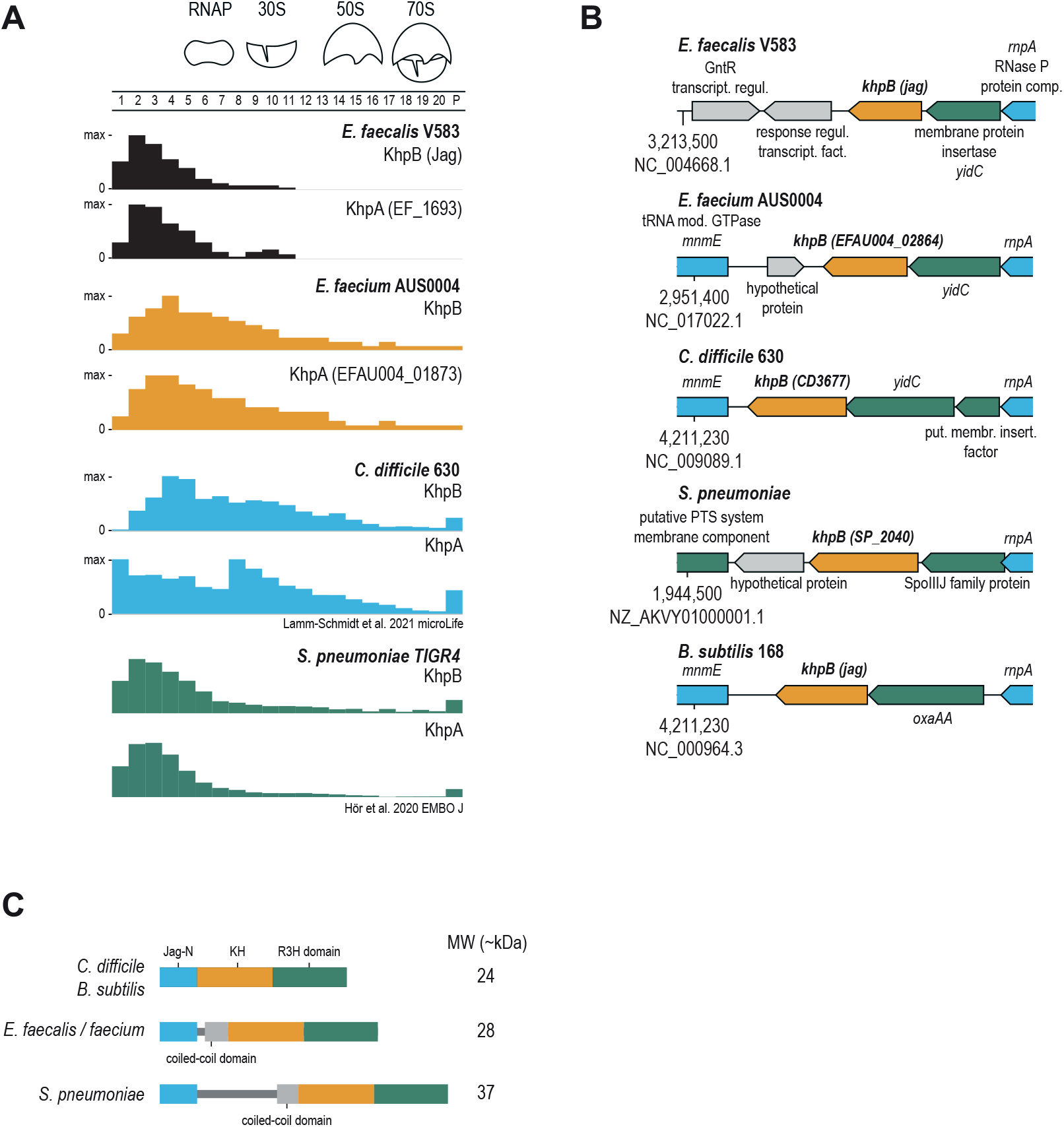
KhpA/B is a conserved RNA-binding protein. (A) Sedimentation profiles of KhpA and KhpB in *E. faecalis*, *E. faecium, C. difficile* and *S. pneumoniae*. (B) Genetic locus of KhpB in different bacterial species. The locus also encodes the RNase P protein component (rnpA) and tRNA modification GTPase (mmE) (C) Domain organization of KhpB in different bacterial species. KhpB can vary in length due to the flexible linker region between the Jag N-terminal domain and the C-terminal KH and R3H domains that are annotated as RNA-binding domains.

KhpA and KhpB behaved similarly in *E. faecalis* and *E. faecium*, with both proteins showing up primarily in the LMW and tailing towards HMW fractions, indicating that they may exist in a free form as well as in a complex, presumably with cellular transcripts (Fig. 5A). However, *E. faecium* KhpA and KhpB showed broader distribution than did their counterparts in *E. faecalis*, which had a similar sedimentation profile to *S. pneumoniae* KhpA and KhpB (Hör et al., 2020b) (Fig. 5A). By contrast, *E. faecium* KhpA and KhpB appeared more similar to Grad-seq profiles obtained for *C. difficile* KhpA and KhpB (Lamm-Schmidt et al., 2021). All these profiles, despite their distinctions, are compatible with the expectation that KhpA and KhpB form RNA-protein complexes.

Comparative analysis of *khpB* genes across Gram-positive bacteria (*C. difficile*, *S. pneumoniae* and *B. subtilis*) suggested a potential conservation of protein function (Fig. 5B). The *khpB* gene lies in close proximity to *yidC* (called *spoIIIJ* in *B. subtilis*), which encodes a protein translocase; *rnpA*, which encodes the RNA component of RNase P; and *mnmE*, which encodes a tRNA modifying GTPase. One characteristic feature of KhpB proteins is their shared domain architecture showing conserved Jag-N, KH and R3H RNA-binding domains (Olejniczak et al., 2021) (Fig. S4A). KhpA proteins, on the other hand, only carry the conserved KH domain. A phylogenetic analysis based on sequence similarity resulted in distinct leaves for KhpA and KhpB, consistent with their overall differences in terms of size and domains (Fig. S4B). It has been noted before that the linker region between the N-terminal Jag-N and KH-R3H domains varies between species (Olejniczak et al., 2021) (Fig. 5C). Notably, in enterococci this linker region harbours a unique coiled-coil domain that could potentially play a structural role and diversify the RNA targetome compared to *Clostridium* and *Streptococcus*.

### A glimpse at the potential cellular RNA targets of KhpB

Based on the interesting sedimentation profile (Fig. 5A) and genetic locus (Fig. 5B), the high sequence conservation (Fig. S4B-C) and unique linker region (Fig. 5C) of KhpB, we sought to obtain a preliminary view of the cellular interactome of this putative RBP. Of the two species in question, we selected *E. faecalis* for further analysis. In order to generate KhpB-specific antisera, we produced the KhpB protein with a C-terminal 3C-cleavage site and a His-tag by heterologous expression in *E. coli*. This recombinant KhpB protein was purified via immobilized nickel affinity chromatography, tag cleavage, and anion exchange chromatography (Figs. 6A, S4C-D) and was used to raise two polyclonal antisera in rabbit that specifically detected KhpB by western blot (Fig. 6B). Of note, raising a KhpB-specific antibody for immunoprecipitaion circumvents exogenous expression of an epitope-tagged KhpB protein construct, as previously done for *S. pneumoniae* or *C. difficile* (Lamm-Schmidt et al., 2021; Zheng et al., 2017), and allows the capture of native KhpB at physiological concentration.

**Figure 6.**
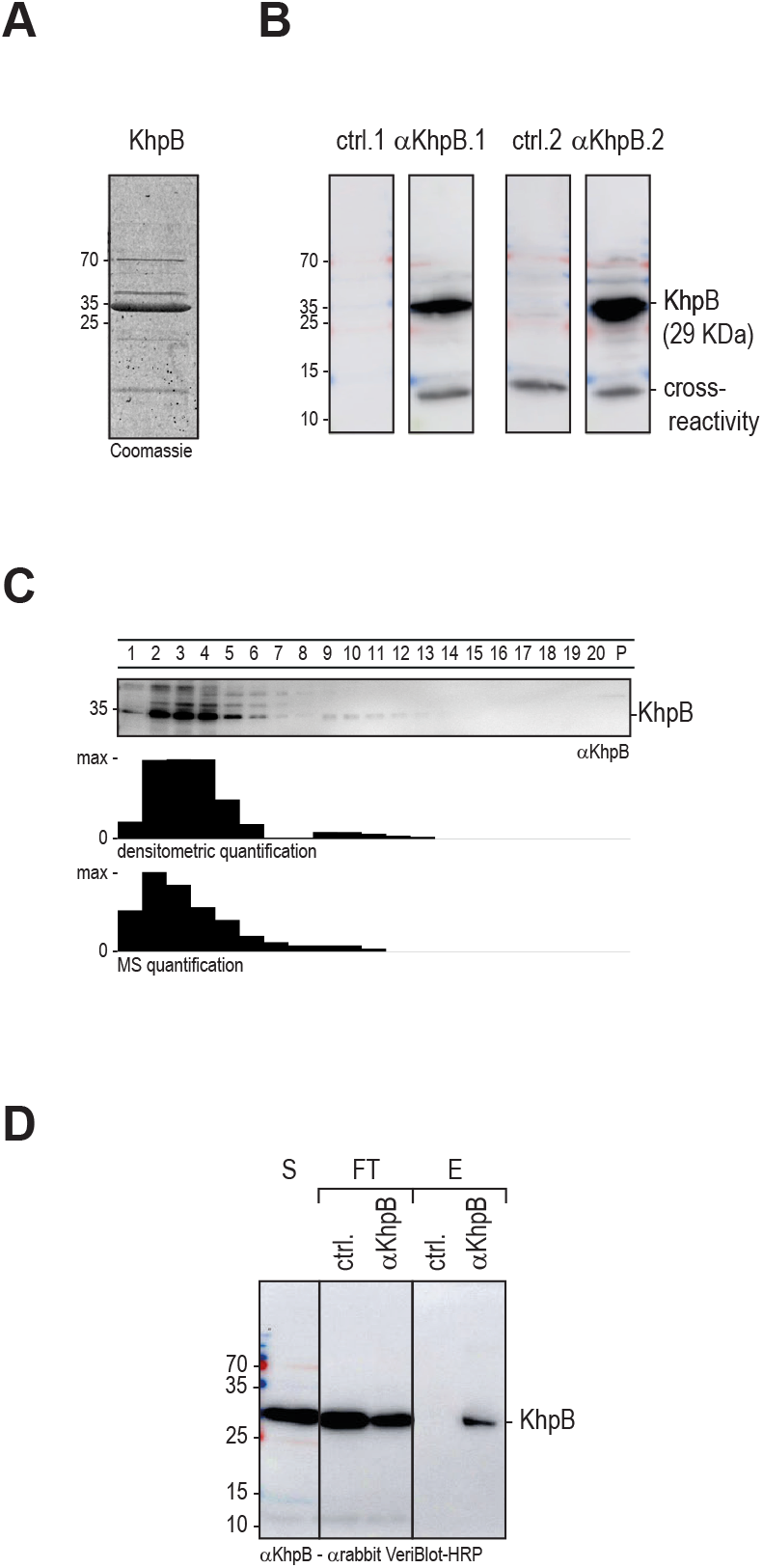
Antibody generation and immunoprecipitation of *E. faecalis* KhpB. (A) SDS-PAGE of purified KhpB protein stained by Coomassi. (B) Western blot of an *E. faeacalis* lysate probed with the two pre-immune sera (ctrl.; left) and the two KhpB anti-sera (right). C) (Top) The sedimentation profile of KhpB detected by western blot analysis. (Middle) Densitometric quantification of the western blot. (Bottom) MS-quantification of the Grad-seq data. (D) KhpB was immunoprecipitated from *E. faecalis* lysate, eluted by organic phase extraction and detected by western blot analysis. S, Supernatant; FT, flow-through; E, eluate; ctrl., pre-immune serum; αKhpB, KhpB anti-serum.

With the α-KhpB sera in hand, we first validated the Grad-seq based sedimentation profile of KhpB by immunoblotting (Fig. 6C). A western blot analysis confirmed the predicted enrichment in LMW fractions 2 to 6 and a faint signal in HMW fractions 9 to 11. Next, we performed RNA immunoprecipitation followed by RNA-seq (RIP-seq) using *E. faecalis* cellular lysates obtained at late exponential phase (OD_600_=2). The experiment was performed twice using one of the two independent α-KhpB antisera per IP. Pre-immune sera were used as a negative control (ctrl., Fig. 6D, Table S5). With the threshold that we applied (log_2_ fold-change >2), determined by deseq2; p-value <0.05 (Wald test)), we recovered cDNA reads for 54 RNAs, showing a nearly even distribution across coding, non-coding, and untranslated regions. A quantitative distribution based on RNA ligand abundance indicates relative enrichment of tRNAs, sRNAs and 3’UTRs, with tRNAs being most enriched. (Fig. 7). KhpB binding seems specific to certain tRNAs; others, despite being encoded in the same genomic locus, showed no enrichment in the IP fraction (Fig. 8).

**Figure 7.**
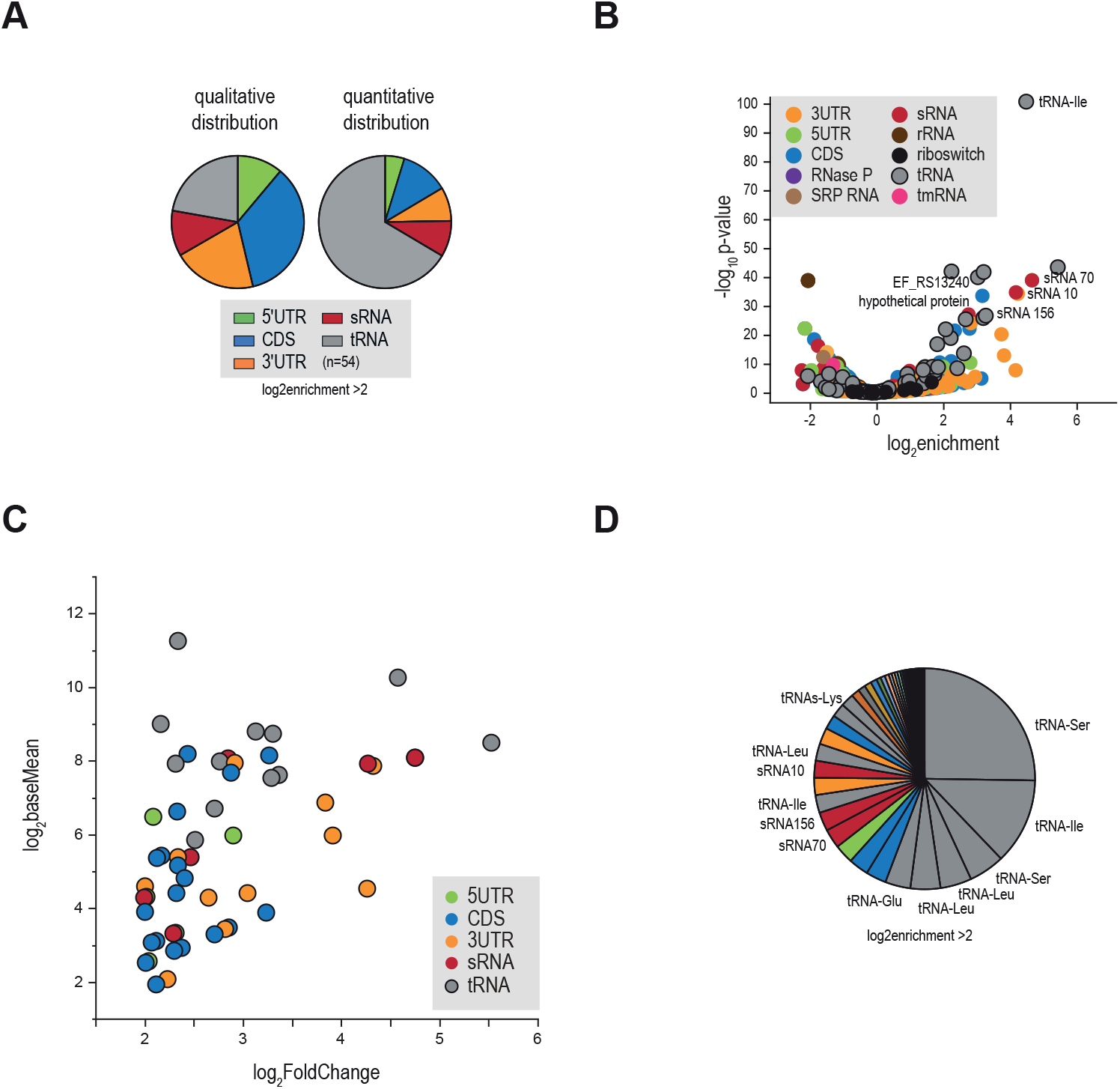
KhpB targets tRNA, sRNAs, and 3’UTRs. (A) Qualitative (left, based on different transcript types) and quantitative (right, based on RNA ligand abundance) distribution of RNAs found enriched in the KhpB immunoprecipitate (log2FC>2). (B) Volcano plot of RNAs associated with KhpB, classified by categories. (C-D) Top enriched RNA candidates were tRNAs, sRNA70, sRNA156 and a number of 3’UTR regions. Reads from two experiments using two independently generated antisera against KhpB were merged by geometrical averaging and the p-values were calculated by the Wald test in deseq2.

**Figure 8.**
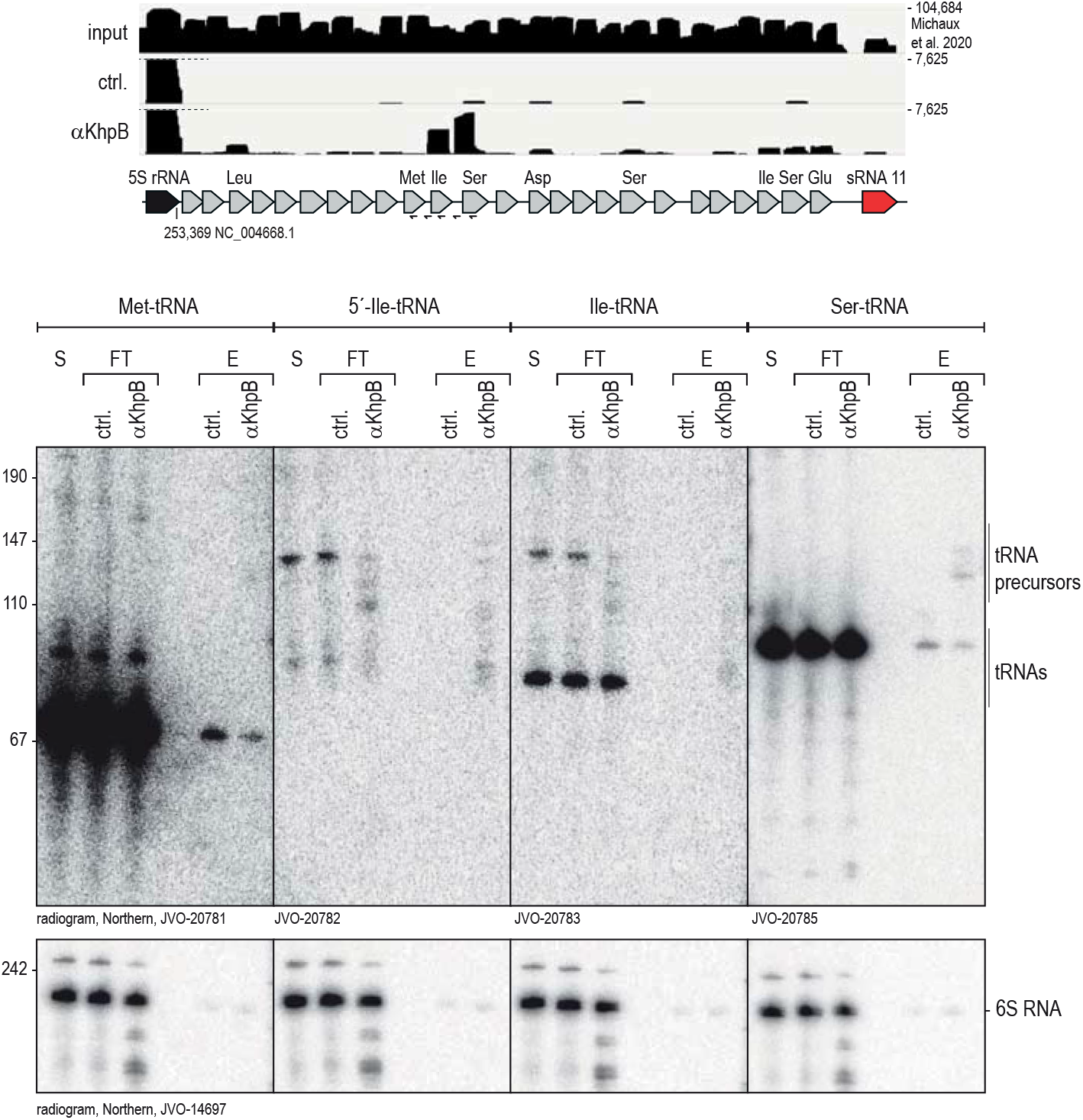
KhpB tRNA target validation. (Top) Read coverage plots of a genomic tRNA locus from *E. faecalis* RNA-seq data (Michaux et al., 2020), the pre-immune serum control (ctrl.) and the KhpB antiserum (αKhpB) pull-down. (Bottom) Northern blot analysis of the immunoprecipitation experiment probed for the indicated tRNAs. S, supernatant; FT, flow-through; E, eluate; JVO, name of the oligoprobes (Table S3).

The limited target suite observed here is in stark contrast to RIP-seq results for KhpB in *C. difficile*, where the IP enriched thousands of different transcripts, including many sRNAs (Lamm-Schmidt et al., 2021). Seeking to validate our RIP-seq data, we probed northern blots of the KhpB immunoprecipitates for tRNAIle and tRNASer and included tRNAMet as a negative control that was not recovered in RIP-seq. tRNA^Met^ was detected in precipitates using both the control and αKhpB immune serum, indicating an non specific interaction. tRNA^Ile^, however, was only detected upon precipitation with the α-KhpB immune serum (Fig. 8). Although the signals in the eluate fractions were generally weak, we did detect bands that correspond to the mature tRNA and to tRNA precursors using probes binding either the 5’UTR of tRNA^Ile^ or within tRNA^Ile^. Similar results were obtained for tRNA^Ser^. Importantly, the read coverage of the tRNA transcripts recovered by RIP-seq extended past the mature 5’end into regions that are known to be processed by RNase P (Klemm et al., 2016), hinting at a potential role of KhpB in tRNA processing. Intriguingly, this would be in line with the genomic location of the *khpB* gene proximal to the genes of the RNase P protein component (*rnpA*) and a tRNA modifying GTPase (*mnmE*) (Fig. 5B).

Our RIP-seq data also predicted KhpB to interact with sRNAs and UTRs (Fig. 7). To validate the interaction between KhpB and the three most enriched sRNAs, northern blots were probed for sRNA70; sRNA10, a putative lysin riboswitch-derived sRNA; and sRNA156, which shares sequence homology with the RNA of a type I toxin-antitoxin module (Fig. 9A-C). For all three, a signal was detected in the RNA extracted after elution with the α-KhpB antisera whereas no signal was detected in the control elution. Interestingly, sRNA156 was the only full-sized sRNA detected; the bands observed for sRNA70 and sRNA10 were shorter compared to the respective sRNA annotation (Michaux et al., 2020) and the signals in the supernatant and flow-through. This observation is another hint at a possible role of KhpB in RNA processing, as is their in-gradient distribution according to the high-throughput Grad-seq data (Fig. 9D). Similarly to KhpB, these sRNAs all occur primarily in fractions 1 to 5.

**Figure 9.**
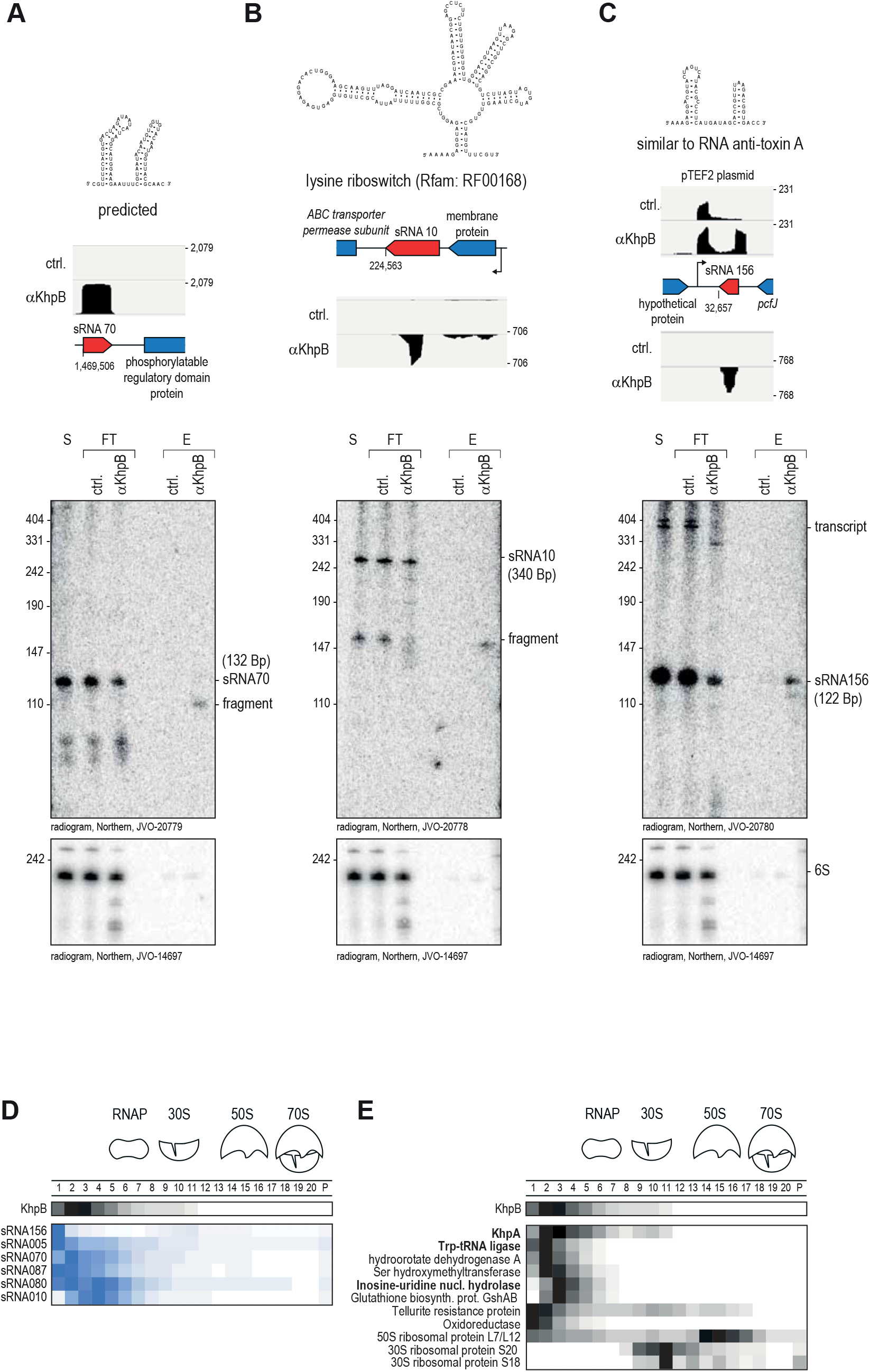
KhpB sRNA target validation. (A-C) Secondary structure predictions, read coverage plots and northern blot validation of sRNAs highly enriched in the KhpB interactome. S, supernatant; FT, flow-through; E, eluate. (D) Comparison between the sedimentation profile of KhpB and the sedimentation profile of some of the most enriched sRNAs of the interactome. (E) Sedimentation profiles of proteins that co-immunoprecipitated with KhpB.

To obtain a first glimpse at potential protein interactors, we also analyzed the KhpB immunoprecipiates by MS (Table S6). Based on this protein interactome list, we reinspected the Grad-seq sedimentation profiles of the KhpB interactors (Fig. 9E). KhpB co-precipitated and co-sedimented with KhpA, indicating that KhpA and KhpB form a complex, as suggested previously in *S. pneumoniae* and *C. difficile* (Lamm-Schmidt et al., 2021; Zheng et al., 2017). Interestingly, tryptophan-tRNA ligase and inosine-uridine nucleoside hydrolase, both enzymes known to act on tRNA, showed similar sedimentation profiles as well.

Altogether, the predicted RNA-targetome of KhpB tentaively suggests interactions with sRNAs, UTRs and tRNA. A potenial functional role for KhpB, e.g., in the processing of these RNAs, and its mode of action will require further investigation.

### A user friendly Gradient browser

We have facilitated access to the enterococcal Grad-seq data via an interactive online browser (Fig. 10). The browser allows the user to search RNA and protein sedimentation profiles for either *E. faecalis* or *E. faecium* (https://resources.helmholtz-hiri.de/gradseqef/) and to manually dissect the profiles for positions and tailing. In addition, candidates belonging to a similar cluster of sedimentation profiles can be displayed automatically. A comparison is possible by adding items to a list, and visualization is provided via bars, plots, and heatmaps. The user can therefore compare protein and RNA sedimentation profiles in order to identify correlated sedimentation and obtain an indication of stable complexes.

**Figure 10.**
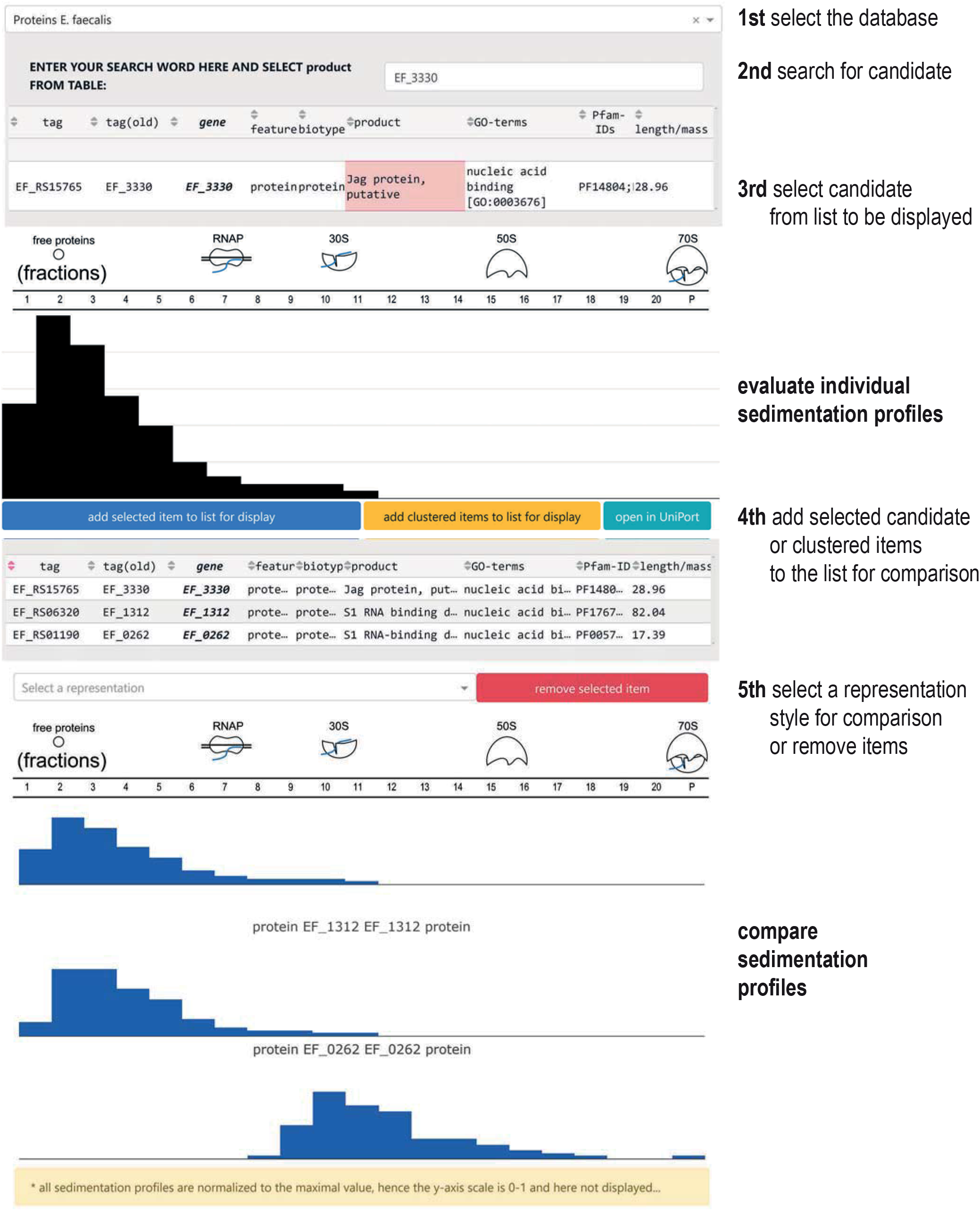
Genome browser. Screen capture illustrated the sedimentation profile browser for *E. faecalis* with a selection of RNA-binding proteins. In the browser both datasets of *E. faecalis* and *E. faecium* can be accessed and used as follows: 1. Selection of the database of interest (*E. faecalis* or *E. faecium)*. 2. Search for the protein or transcript of interest by name. (3) Select the candidate of interest. (4) Display of individual sedimentation profiles. (4) Add candidates to the list for further comparison. (5) Select the style of visualisation for comparison. (6) Compare sedimentation profiles.

## DISCUSSION

Grad-seq is a powerful method to study native RNA-protein complexes. Its use was first demonstrated through the identification of ProQ as a new global RBP in *Salmonella* (Smirnov et al., 2016). Subsequent Grad-seq studies have put the spotlight on additional RBPs, such as KhpB in *C. difficile* (Lamm-Schmidt et al., 2021), or revealed a coding capacity of RNAs previously thought to be noncoding, as demonstrated in *E. coli* for the sRNA RyeG through very similar in-gradient profiles of the sRNA RyeG and the 30S ribosomal subunit (Hör et al., 2020a). Here, we have used this approach to provide a global resource for the identification of stable RNA and protein complexes in the human pathogens *E. faecalis* and *E. faecium*.

Several sRNAs of *E. faecalis* and *E. faecium* have previously been implicated in stress responses and virulence phenotypes (Fouquier d’Hérouel et al., 2011; Innocenti et al., 2015; Michaux et al., 2014; Shioya et al., 2011; Sinel et al., 2017), suggesting their importance in post-transcriptional gene regulation in these two bacterial species. Recently, we systematically annotated sRNAs in *E. faecalis* and *E. faecium* based on dRNA-seq data (Michaux et al., 2020), but this approach works less well for the discovery of UTR-derived and processed sRNAs that lack primary transcript ends. Therefore, UTR-derived sRNAs might have been missed in our previous annotation. Using our Grad-seq dataset, we were able to re-assign UTRs that showed atypical sedimentation profiles as non-coding RNAs, leading to the identification of multiple additional enterococcal sRNA candidates that are generated from mRNA 5’ or 3’ UTRs. We are still at the beginning of understanding to what extent mRNA-derived sRNAs differ in regulatory scope and function from canonical sRNAs. Nevertheless, several such UTR-derived sRNAs have been characterized in other species, often revealing unexpected, conserved functions in diverse cellular processes (Adams and Storz, 2020; Ponath et al., 2022). Identification of the cellular targets of the UTR-derived sRNAs that we predict based on our Grad-seq data will be the next step for future studies using methods of the bacterial sRNA tool kit (Hör et al., 2018; Jagodnik et al., 2017).

Interestingly, while sRNAs are enriched in the LMW fractions, they also show a broader distribution in HMW fractions (Gerovac et al., 2020). This implies that they exist both in a free form and in larger complexes, indicating that *Enterococcus* sRNAs, like the sRNAs in other bacterial species, interact with other RNAs or with proteins. Like most Gram-positive bacteria, neither *E. faecalis* nor *E. faecium* encode any of the well-known sRNA chaperones such as Hfq and ProQ. Regarding putative new sRNA-related proteins, our Grad-seq revealed similar sedimentation between KhpA, KhpB, and many sRNAs. In addition, both proteins were previously shown to interact with sRNAs in *S. pneumoniae* and *C. difficile* (Lamm-Schmidt et al., 2021; Zheng et al., 2017). Indeed, our preliminary RIP-seq data predict *E. faecalis* KhpB to interact with tRNAs, sRNAs, and 3’UTRs. While sRNAs are common interaction partners of KhpB in all bacterial species studied so far, *C. difficile* KhpB was not seen to interact with 3’UTRs. This may be because *C. difficile* possesses Hfq and ProQ, both known to bind to the 5’ and 3’ ends of mRNAs. *S. pneumoniae* does not encode Hfq, and the KhpB interactome featured UTRs, like the *Enterococcus* KhpB interactome. This might imply that in bacterial species that lack Hfq and ProQ, UTRs are targeted by KhpB.

Our data also suggest that KhpB might have a role in tRNA and sRNA processing, but future work is required to establish such a function. Curiously, both tRNA^Tyr^ and tRNA^Pro^ were also found to be RNA ligands of KhpB in *S. pneumoniae* (Zheng et al., 2017), but tRNAs do not interact with KhpB in *C. difficile*, despite the much broader interactome of *C. difficile* KhpB (Lamm-Schmidt et al., 2021). At any rate, our RIP-seq data are encouraging and could in future studies be followed up using more stringent RNA-protein interactome methods, foremost those that include UV crosslinking (Holmqvist et al., 2016; Tree et al., 2014) to form native bonds between an RBP and its RNA targets. One major advantage of these methods is that they enable homing in on the transcript region recognized by the RBP of interest; this extra positional information will be important to validate the putative role of KhpB in guiding RNA processing of noncoding transcripts. Other open questions remain: for instance, we show that KhpB interacts with KhpA, as previously reported (Winther et al., 2019; Zheng et al., 2017), but whether both proteins interact with RNA cooperatively will need to be further examined. Whether or not KhpB is in fact invovled in sRNA pairing with target mRNAs is still unknown as well. Further analysis will be requried to dissect the functional role and mode of action of KhpB in enterococci.

Overall, we believe that our Grad-seq data provides a powerful resource and ample departure points to identify RNA-protein complexes of *E. faecalis* and *E. faecium*. We are confident that this data will aid in the investigation of post-transcriptional regulation in these opportunistic pathogens, an area of molecular microbiology that is still in its infancy. Of note, we recently expanded the molecular toolkit for the global identification of RNA-protein interactions by developing SEC-seq, a method that couples size exclusion chromatography with RNA-sequencing and mass-spectrometry (Chihara et al., 2022). It is based on a similar concept as Grad-seq, but shows improved resolution in the LMW range, a size range that is particularly relevant for the analysis of regulatory RNA-protein complexes in bacteria. Therefore, it would be interesting in the future to build on our Grad-seq data for *E. faecalis* and *E. faecium* by performing SEC-seq on these species.

---

**Figure S1.**
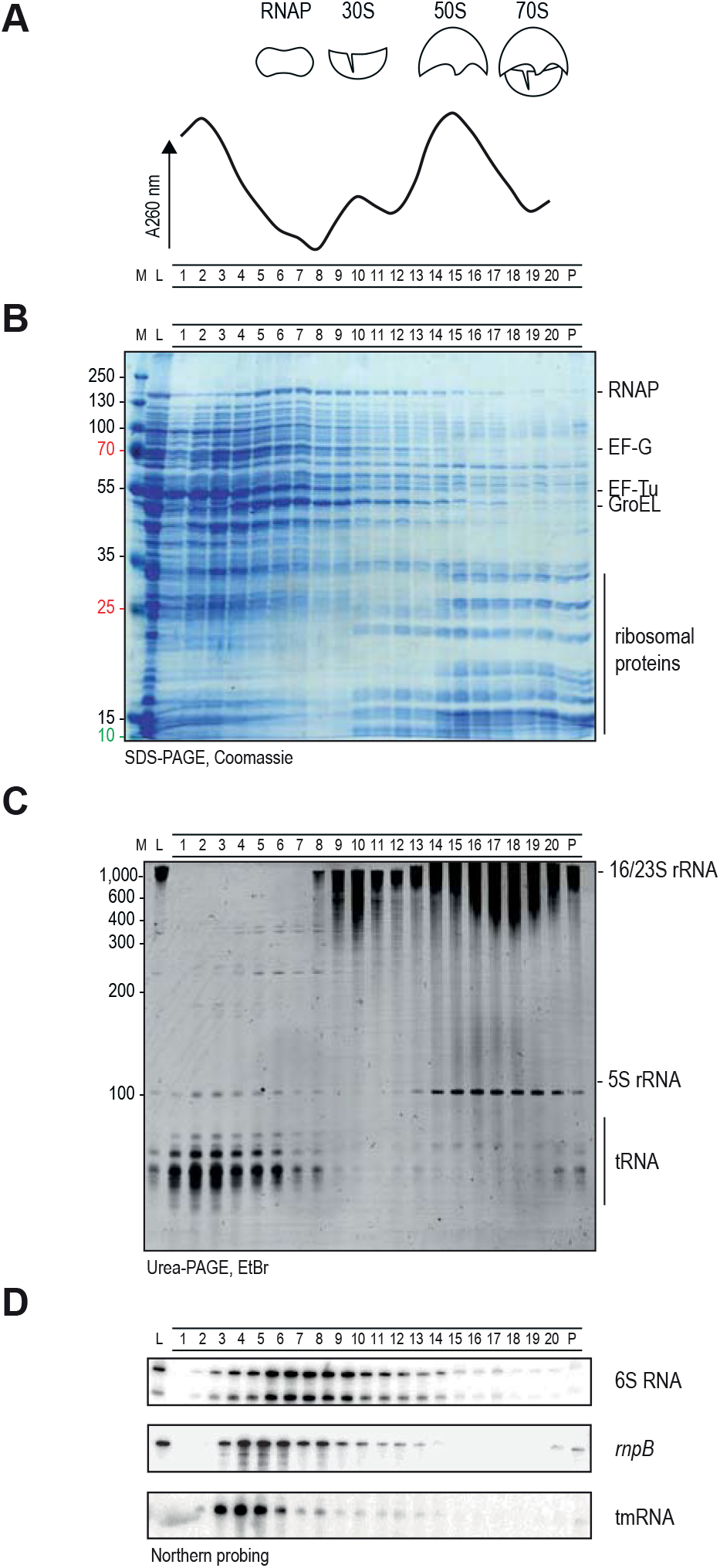
Grad-seq in *E. faecium*. (A) Absorption profile at 260nm of the gradient fractions. (B) Fractions were run on an SDS-PAGE and abundant proteins were detected by Coomassie staining. (C) Fractions were run on a Urea-PAGE and abundant RNAs were detected by ethidium bromide staining. (D) Northern blots were probed for the indicated RNAs to assess their sedimentation profile.

**Figure S2.**
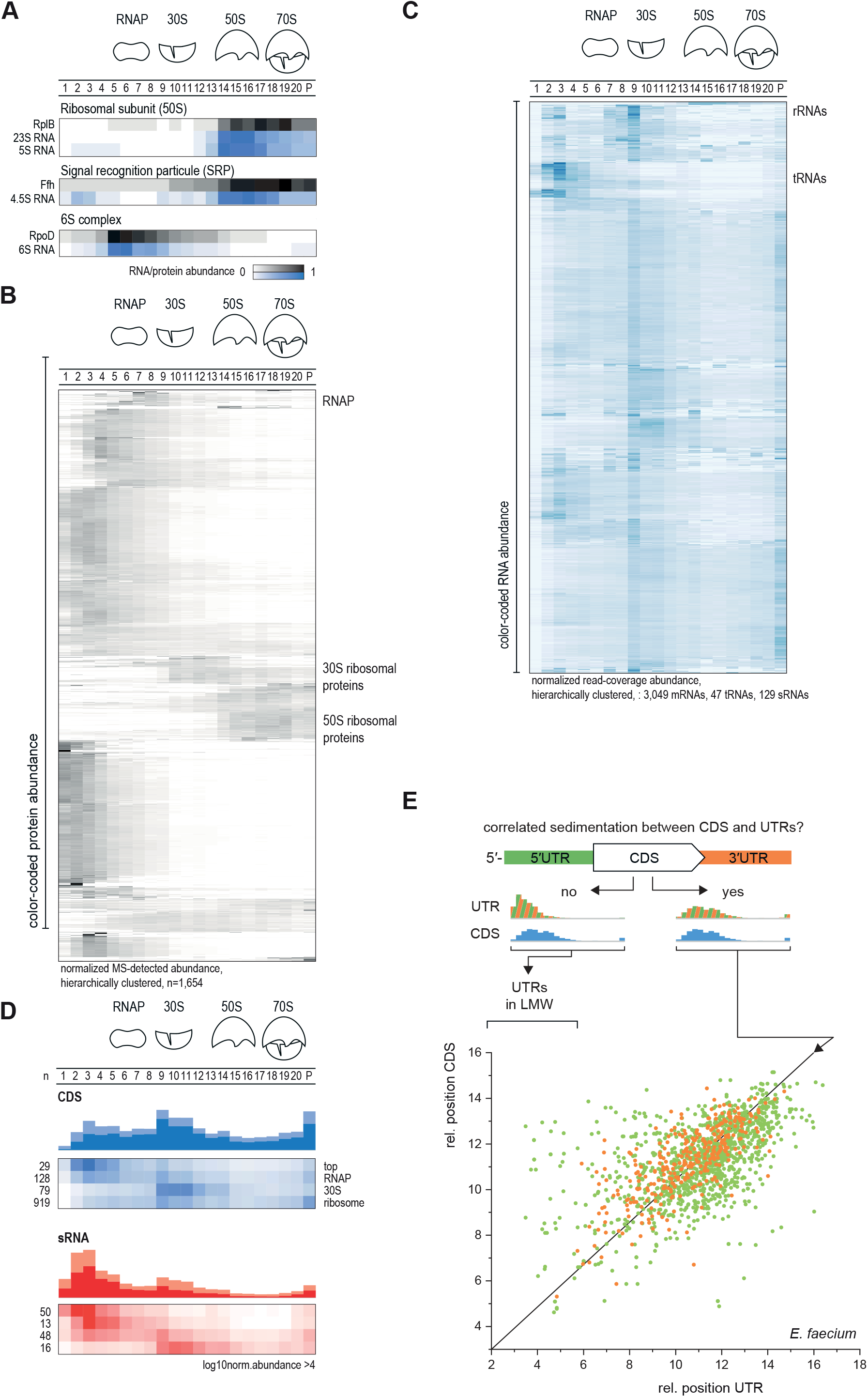
Sedimentation profiles in Grad-seq for *E. faecium*. (A) Sedimentation profiles of known RNA-protein complexes, specifically the ribosomal 50S subunit complex, the signal recognition particle (SRP) and the 6S complex. (B) Heatmaps based on unsupervised clustering of the sedimentation profiles of *E. faecalis* proteins. (C) Heatmaps based on unsupervised clustering of the sedimentation profiles of *E. faecalis* RNA. RNA sedimentation profile patterns can be detected, for example ribosomal RNAs and tRNAs. (D) Merged profiles of the sRNAs and CDS sedimentation profiles. (E) Comparison of the relative positions of CDSs and their UTRs in the sedimentation gradient. Outliers indicate a different sedimentation between the UTR and the CDS.

**Figure S3.**
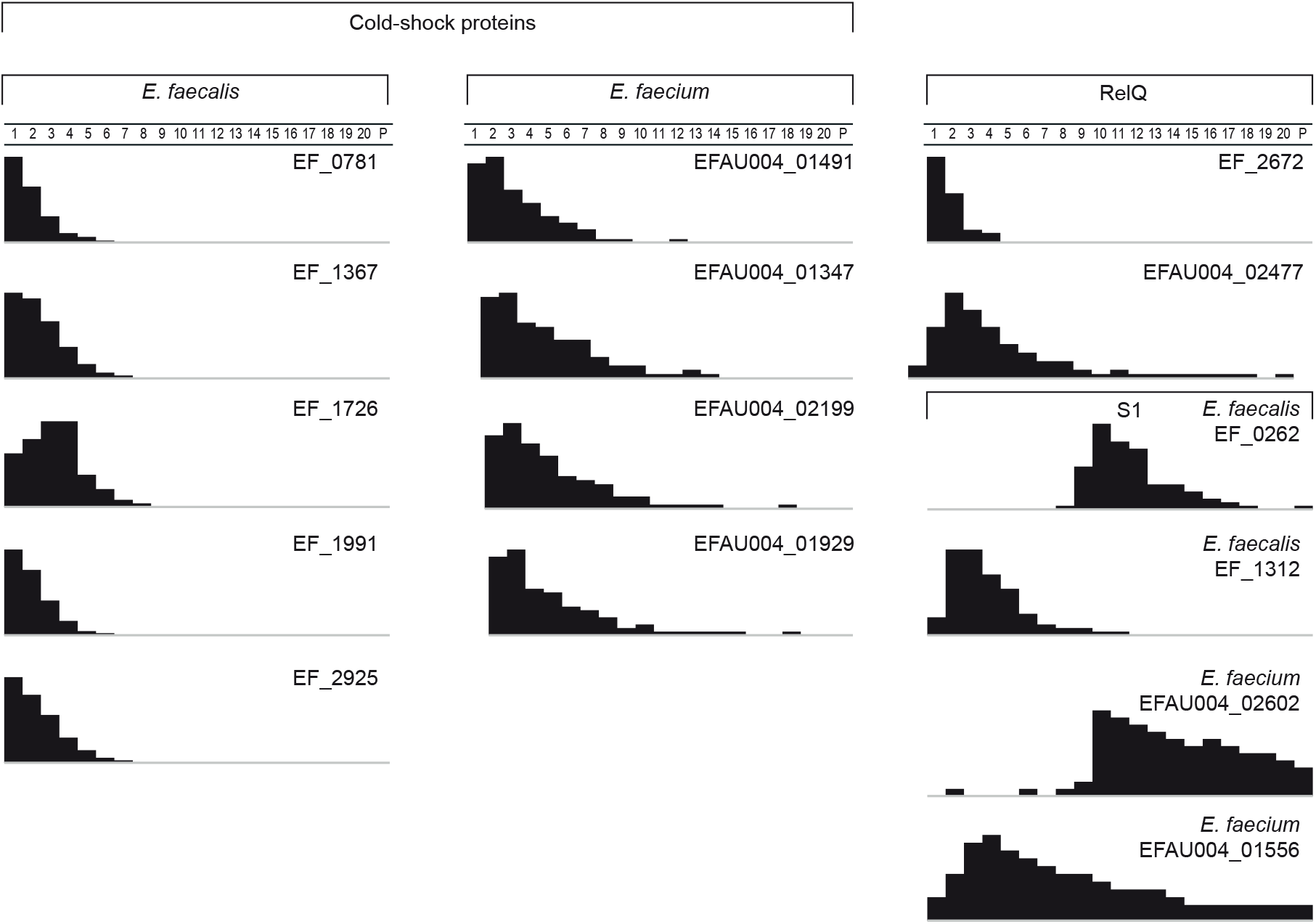
Grad-seq sedimentation profiles of known RNA-binding proteins. Depicted are the profiles of the cold-shock proteins, RelQ and S1 proteins for both *E. faecalis* and *E. faecium*.

**Figure S4.**
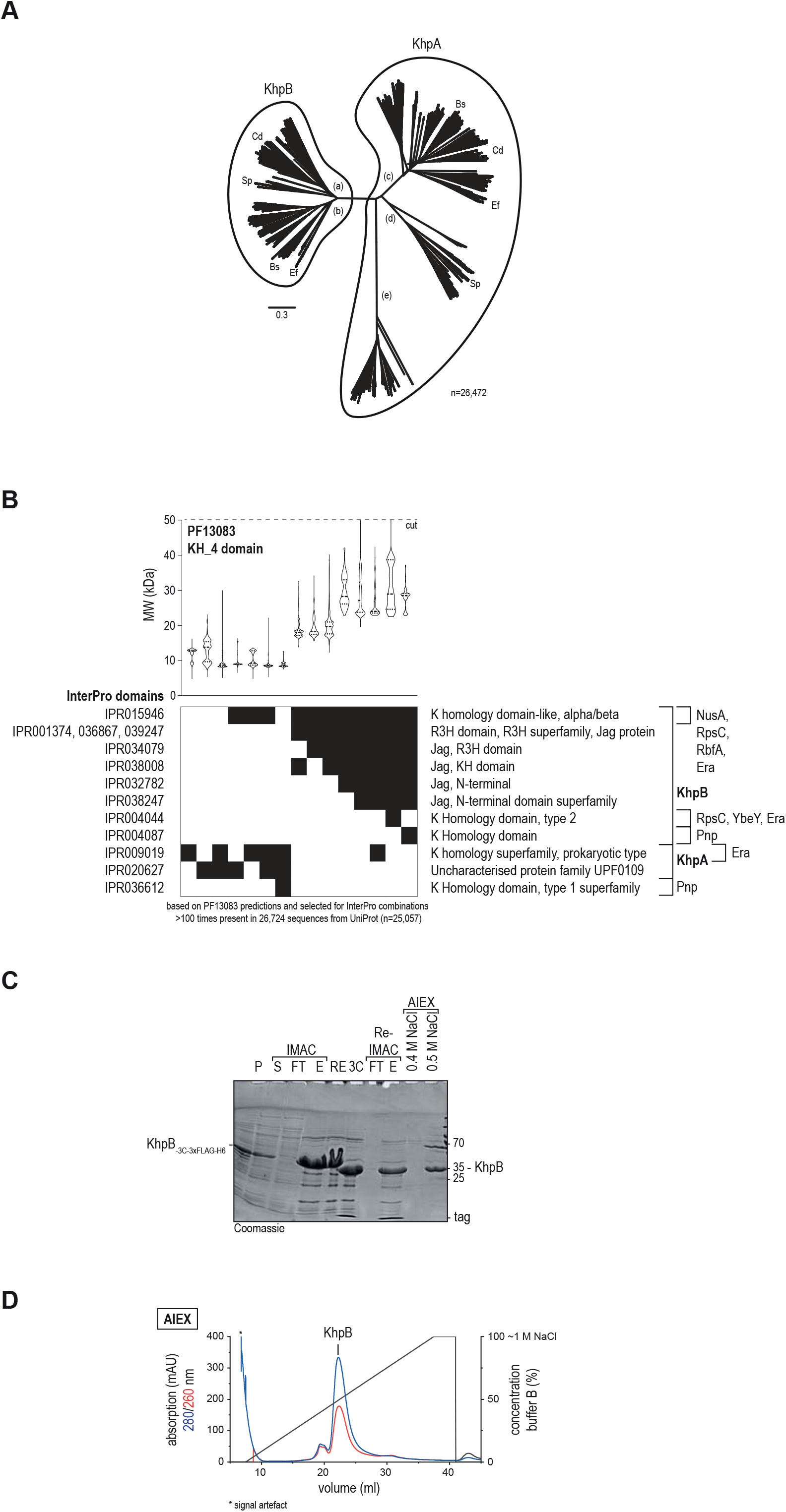
Conservation of KhpB and protein purification. (A) KhpA and KhpB conservation tree by sequence homology based on 26,472 sequences from UniProt. (B) KH-domain protein sub classification by InterPro annotated domains. (C) KhpB protein purification via immobilized metal affinity chromatography (IMAC), 3C cleavage, reverse IMAC, and anion exchange chromatography. KhpB was eluted at 0.5 M NaCl and was concentrated and rebuffered. P, pellet; S, supernatant; FT, flow-through; E, elution; RE, rebuffered sample; 3C, 3C-cleaved sample. (D) Anion exchange chromatography (AIEX) elution profile of KhpB. * signal artifact.

**Table S1.** Mass-spectrometry results obtained from the proteins extracted on the 20 fractions of the glycerol gradient performed for *E. faecalis* and *E. faecium*, normalized and classified per fraction.

**Table S2.** RNA-seq results obtained from the RNA extracted on the 20 fractions of the glycerol gradient performed for *E. faecalis* and *E. faecium*, normalized and classified per fraction.

**Table S3.** Combined information on the material used in this study: oligonucleotide sequences, the plasmids used for the protein expression and purification, the strains, the antibody specification, the software and codes used to analyze the data and the identifiers for data deposition in GEO and PRIDE.

**Table S4.** Sedimentation of the CDS and corresponding 5’ and 3’UTRs (related to Fig. 3).

**Table S5.** RNA interaction partners of KhpB. For each RNA, name, RNA classification, enrichment fold and p-value are indicated. The RIP-seq experiment was conducted with two independently generated antibody sera raised against KhpB. Deseq2 was used to geometrically average and normalize read counts per sample and to calculate the p-value using the Wald test.

**Table S6.** Protein interaction partners of KhpB.

## MATERIALS AND METHODS

### Bacterial growth and culture conditions

Bacteria, *E. faecalis* V583 and data *E. faecium* AUS004, gift from Pr. Jean-Christophe Giard, were grown at 37°C on M17 agar plates (Oxoid) supplemented with 0.5% glucose. Single colonies were inoculated each time liquid cultures were needed. 8.3ml of M17 medium supplemented with 0.5% glucose were inoculated in a 25ml glass tube, in order to keep the final proportions of one third of media and two thirds oxygen. Cultures were grown overnight at 37°C and back diluted 1:100 into fresh media and grown without agitation to late logarithmic/early stationary phase at an OD_600_ of 2.0. For the glycerol gradient sample preparation, 200 OD_600_ of bacterial culture was used for each strain. 1ml of the overnight culture was diluted in 100ml of GM17 and grown to OD_600_ of 2.

Once the desired OD was reached, the cultures were decanted into pre-chilled 50ml falcons and incubated on ice for 15min with rotation of the tubes every 3-5min to ensure fast and uniform cooling. The cells were harvested by centrifugation at 4000g at 4°C for 20min in 50ml falcon tubes. Pellets were pooled in order to have one tube per strain. Three washes with 25ml of ice-cold 1xTBS were applied and the pellets were resuspended in a final 1ml 1xTBS, transferred into a 2ml tube and centrifuged at 13000 rpm, 4°C, during 2 min. After removing all the TBS, pellets were resuspended in 500 μl of lysis buffer (20 mM Tris-HCl, pH 7.5, 150 mM KCl, 1 mM MgCl_2_, 1 mM DTT, 1 mM PMSF, 0.2 % Triton X 100, 20 U/ml DNase I, 200 U/ml RNase-inhibitor). An equal volume of 0.1 mm glass beads was added, and cells were vortexed for 30 sec followed by 15 sec cooling on ice. This lysis step was repeated 10 times.

The lysate was then cleared by centrifugation at 13,000 rpm, 4°C during 10 min. Without disturbing the pellet, all supernatant was collected. A couple of aliquots were saved (Protein: 20 μl lysate + 20 μl of 5x Laemmli buffer; RNA: 10μl lysate + 1 ml Trizol in 2 ml SafeSeal tube) for subsequent analysis of proteins and RNAs. 200 μl of the very top part of the gradient was removed and 200 μl of the cleared lysate was gently placed. The gradient used here was a linear 10-40% (w/v) glycerol gradient. To prepare the gradient, two solutions were made. The 10% (w/v) glycerol solution (20 mM Tris-HCl, pH 7.5, 150 mM KCl, 1 mM MgCl_2_, 1 mM DTT, 1 mM PMSF, 0.2 % Triton X 100, 10 % (w/v) glycerol) and the 40% (w/v) glycerol solution (20 mM Tris-HCl, pH 7.5, 150 mM KCl, 1 mM MgCl_2_, 1 mM DTT, 1 mM PMSF, 0.2 % Triton X 100, 40 % (w/v) glycerol). Clean ultracentrifugation tubes were divided into two. The first half of the tube was filled up with the 10% solution and using a 70 mm injection needle and a syringe, the 40% solution was then placed underneath the 10% solution until the interphase reached the middle of the tube. The loaded gradients were centrifuged at 100,000 g (23,700 rpm) for 17h at 4°C in a precooled ultracentrifuge. The gradients were then manually fractionated into 590 μl fractions. Following this method, 20 fractions and the pellet were collected for each gradient (separately for *E. faecalis* and *E. faecium*).

### Protein / mass spectrometry sample preparation

From each fraction, 90 μl was mixing with 30 μl of 5x Laemmli loading buffer. The fractions were analyzed in 12% SDS-Page standard gel followed by Coomassie staining (20 μl for the fractions, 3 μl for the lysate and 10 μl for the pellet). KhpB was detected by the αKhpB antibody (2, SY0917 (MA11010) SABC-Serum) (rabbit, 1:10,000) and a secondary anti-rabbit-HRP (goat) conjugate antibody (1:10,000, Sigma-Aldrich, #31460). The rest of the samples was prepared for mass spectrometric analysis as previously described (Hör et al., 2020b). Briefly, samples were homogenized using ultrasound. Debris were later removed by centrifugation and 20 μl of the cleared protein sample were spiked-in with 10 μl of UPS2 spike-in (Sigma-Aldrich) diluted in 250 μl of 1.25x protein loading buffer. The samples were then reduced in 50mM DTT, 10 min at 70°C and alkylated with 120 mM iodoacetamide for 20 min at room temperature in the dark. The proteins were precipitated using 4 times their volume of acetone overnight at −20 °C, and later on washed with acetone and dissolved in 50 μl of 8 M urea, 100mM ammonium bicarbonate. Protein digestion into peptides was performed using Lys-C (Wako) for 2h at 30°C following by overnight digestion by trypsin. Peptides were eluted with 60% acetonitrile/0.3% formic acid and stored at −20°C until LC-MS7MS analysis.

### RNA / RNA-seq sample preparation

From each collected fraction (minus the 90 μl taken for protein preparation), 50 μl of 10% SDS was added (or 25 μl for the pellet as the volume was expectedly lower in the sample). After shaking by hand vigorously for about 20 sec, 600 μl of acidic P:C:I was added to each fraction (300 μl for the pellet). 400 μl of chloroform was added also to the Trizol-dissolved lysate and shaken by hand for about 10 seconds. All the samples were then vortexed for 30 sec and let rest for 5 min at room temperature. The samples were centrifuged at 13000 rom at 4°C for 15 min. The aqueous phases were collected in 2ml tubes and 1 μl of Glycoblue, together with 1.4 ml ice-cold 30:1 EtOH:3 M NaOAc pH 6.5 were added. Tubes were put at −80°C for 40 min in order to let the RNA precipitate. The samples were then centrifuged at 13000 rpm, 4°C for 30 min, pellets were washed with 350 μl of ice-cold 70% EtOH and centrifuged again for 10 min at 13000 rpm, at 4°C. After air-drying the pellets, 40 μl nuclease-free water was added. DNase treatment was performed. For one entire gradient, a master mix was prepared: 115 μl of DNase I buffer with MgCl_2_, 11.5 μl RNase-inhibitor, 92 μl DNase I and 11.5 μl nuclease-free water. RNA were denaturated at 65°C for 5 min prior to treatment. 10 μl of the master mix was added to each 40 μl sample, and the now-treated samples were incubated at 37°C for 45 min. 150 μl of nuclease-free water was then added along with 200 μl of acidic P:C:I. After 5 min of incubation at room temperature, samples were centrifuged at 13000 rpm at 4°C for 15 min, the aqueous phase was placed in the precipitation mix (600 μl ice-cold 30:1 EtOH:3M NaOAc pH 6.5 with 1 μl of Glycoblue), 40 min at −80°C, centrifuged at 13000 rpm 4°C for 30 min, washed with 350 μl 70% EtOH, and centrifuged at 13000 rpm 4°C for 10 min. The pellets were air-dried and dissolved in 35 μl nuclease-free water. 10 μl of each sample was mixed with 10 μl of GLII RNA loading buffer.

For quality control of the RNA, a thick 6% PAA gel with 7M urea was cast with a 25 well comb in order to load the 20 fractions but also the lysate and the pellet. 6 μl of each sample was loaded and run at 300V for about 1h40. The gel was then stained using Ethidium bromide. The same procedure was followed for northern blot preparation. After the run, RNA was transferred onto a Hybond-XL membrane and hybridized overnight at 42°C with γ^32^P-ATP end-labeled oligodeoxynucleotide probes (listed in Table S3). Signals were visualized on a Typhoon FLA 7000 Phosphoimager and quantified with ImageJ (EMBL software publicly available).

For RNA-seq, 5 μl of the gradient samples was diluted in 45 μl nuclease-free water. 10 μl of this dilution was mixed with 10 μl of a 1:100 dilution of the ERCC spike-in mix 2 (Thermo Fisher Scientific) and subjected to library preparation for next-generation sequencing (Vertis Biotechnologie) following the protocol that has been previously described (Hör et al., 2020b).

### RNA-seq and MS analysis

Both analyses were conducted as previously described (Gerovac et al., 2021b). For RNA-sequencing analysis, we used the NCBI reference sequences NC_004668.1, NC_004669.1, NC_004670.1, and NC_004671.1 for *E. faecalis* and NC_017022.1, NC_017023.1, NC_017024.1, and NC_017032.1 for *E. faecium*. Annotation files were used from NCBI. ERCC spike-in reference sequences were also included. RNA-sequencing analysis including read mapping, generation of coverage files, gene quantification, and differential gene expression/level analysis (deseq2) were conducted in READemption 0.4.5 (Förstner et al., 2014). The relative position of an annotated CDS or UTR was determined by the sum of the read fraction per gradient multiplied with the gradient fraction number (P=21), as described previously (Gerovac et al., 2020). Relative positions between CDS and UTRs were correlated and differences of 5 in relative positions indicated independent behavior in gradients.

For protein searches by MS, we used the UniProt proteome UP000001415 for *E. faecalis* and UP000007591 for *E. faecium*.

### KhpB expression and purification

KhpB was cloned for expression into the vector pETM14 (kanamycin resistance) with C-terminal 3C-cleavage site, 3×FLAG-tag, and 6×His-tag. The *khpB* insert was amplified by primers JVO-19671×JVO-19672. The vector backbone was amplified from the template pMiG006 (Gerovac et al., 2020) by JVO-19668×JVO-19669 and JVO-19670 subsequently to add the C-terminal tag. Insert and Backbone were cloned by Gibson assembly. The final plasmid pMiG034 was sequenced for verification by T7 primer.

For expression pMiG-034 was transformed into *E. coli* BL21-CodonPlus (DE3)-RIL cells (Agilent Technologies, JVS-12280, chloramphenicol resistance). Cells were grown in LB to OD_600_ 0.6 at 37°C, cooled to 18 °C and induced with 0.5 mM IPTG overnight. Cells were resuspended in lysis buffer (20 mM Tris/HCl pH 8.0, 1 M NaCl, 0.1 mM EDTA, 4 mM 2ME, 1 mM PMSF) and disrupted by sonication (50% amplitude, 30 s pulsation — 30 s break for 5 min, on ice, Sonopuls HD 3200, TT13 tip, Bandelin). The lysate was cleared at 15,000 × g for 15 min at 4 °C. Supernatant was loaded onto an Ni-NTA column (5 ml, HisTrap HP, Cytiva) and washed with 5 column volumes. Protein was eluted with 200 mM imidazole and desalted (PD10 column, GE) in 3C-cleavage buffer (50 mM Tris/HCl pH 7.5, 150 mM NaCl, 0.1 mM EDTA, 1 mM TCEP). Cleavage of the tag was performed in 2-3 ml with 20 μl 3C protease (1 mg/ml, Sigma-Aldrich, SAE0045) added and incubated over-night at 4°C. The sample was loaded again on an Ni-NTA column for reverse-IMAC. The protein remained bound to the column, and was washed, and eluted again. The sample was diluted with 10 volumes anion exchange buffer A (20 mM HEPES pH6.0, 50 mM NaCl, 0.1 mM EDTA, 4 mM 2ME) and loaded onto an anion exchange column (5 ml, HiTrap Q HP, Cytiva). Protein was eluted in a gradient up to 1 M NaCl. KhpB eluted as a second peak ~0.5 M NaCl. The peak was pooled and concentrated by centrifugal filtration (Amicon Ultra-4, 10 kDa cut-off, Merck) rebuffered in storage buffer (20 mM HEPES/KOH pH 7.5, 150 mM KCl, 1 mM TCEP) and flash frozen in liquid nitrogen. The concentration of KhpB was estimated by absorption measurement at 280 nm using the extinction coefficient of 10,430 mM^−1^cm^−1^. 0.8 mg protein was used for raising two antibodies against KhpB in rabbit (Eurogentec, speedy program).

### RIP-seq

300 μL polyclonal anti-KhpB serum (SY0916 (MA11009) SABC-sérum for experiment 1 and SY0917 (MA11010) for experiment 2), or corresponding pre-immune sera for negative control, was added to 75 μl of prewashed protein A sepharose beads (Sigma-Aldrich) together with 700 μl lysis buffer (20 mM Tris/HCl pH 8.0, 150 mM KCl, 1 mM MgCl_2_, 1 mM DTT) and incubated for 1 h. Beads were washed two times with lysis buffer. Sixty OD of *E. faecalis* cells at OD 2 were resuspended in 1 ml lysis buffer (plus 2 mM PMSF, 10 μl lysozyme (50 mg/ml), 5 μl RNase inhibitor, 5 μl DNase I), and were lysed via FastPrep with lysing matrix E at 6 m/s for 40 s.

The lysate was cleared for 10 min at full speed at 4 °C. Beads were added to the lysate and incubated at 4 °C for 2 h with rotation. Beads were pelleted at 300 × g and washed five times with lysis buffer. Beads were resuspended in 500 μl lysis buffer and RNA was extracted by addition of 1% (w/v) SDS and one volume PCI. The aqueous layer was re-extracted with chloroform and the RNA was precipitated with 0.1 M sodium acetate and 3 volumes of ethanol. RNA was pelleted, dried, and solubilized in water. DNA was digested with DNase I in reaction buffer (Thermo Scientific). RNA was re-extracted by PCI, and the concentration was determined by absorption at 260 nm. Library preparation and sequencing was conducted at Vertis Biotechnologie AG. RNA-sequencing was analysed as described previously. Read quantification between both experiments with independently generated antibodies against KhpB was merged and normalized in deseq2 by geometrical averaging and p-values were calculated using the Wald-test.

The organic layer was precipitated with 10 volumes methanol and LS-MS/MS analyzed for precipitated proteins at the MS core unit (AG Schlosser, Rudolf Virchow Center for Integrative and Translational Bioimaging). RNA extracted from supernatant, the flow-through with either the pre-immune serum or with the anti-KhpB serum and the elution with either the pre-immune serum or the anti-KhpB serum were used to validate KhpB targets by northern blot assays with a similar protocol described in the previous session “RNA /RNA-seq sample preparation”. The probes used against tRNA^Met^, tRNA^Ile^, tRNA^Ser^, the 5’UTR of tRNA^Ile^ and tRNA^Ser^, the sRNA 70, sRNA 10 and sRNA 156 are listed in Table S3.

## Data availability

MS data are accessible at the ProteomeXchange consortium (Deutsch et al., 2017) via the PRIDE partner repository (Perez-Riverol et al., 2015) with the data set identifier PXD031577. Processed data obtained after MaxQuant and sequencing analysis are listed in Table S1 for both species. Grad-seq and KhpB RIP-seq RNA-sequencing raw FASTQ and analysed WIG and TDF coverage files are accessible at Gene Expression Omnibus (GEO) (Edgar et al., 2002) with the accession number GSE196534. The code for the Grad-seq browser is deposited at Zenodo (https://zenodo.org/record/3955585). The Grad-seq browser is online, accessible at https://resources.helmholtz-hiri.de/gradseqef/. READemption 0.4.5 is deposited at Zenodo 1134354 (https://zenodo.org/record/1134354).

## ACKNOWLEDGEMENTS

The authors would like to thank Jean-Christophe Giard for providing the *E. faecalis* and *E. faecium* strains used here, Jens Hör for advice on Grad-seq of Gram-positive bacteria, and members of the Vogel lab for comments, and Anke Sparmann for editing the manuscript. This work was supported by a Gottfried Wilhelm Leibniz Prize awarded to JV (grant DFG Vo875-18).

## CONTRIBUTIONS

CM, MG and JV designed the research. CM, MG, and EEH performed research. CM, MG, EEH, LB, and JV analysed data. CM, MG, EEH and JV wrote the paper with input of all the authors.

